# Triaging of ⍺-helical proteins to the mitochondrial outer membrane by distinct chaperone machinery based on substrate topology

**DOI:** 10.1101/2023.08.16.553624

**Authors:** Gayathri Muthukumar, Taylor A. Stevens, Alison J. Inglis, Theodore K. Esantsi, Reuben A. Saunders, Fabian Schulte, Rebecca M. Voorhees, Alina Guna, Jonathan S. Weissman

## Abstract

Mitochondrial outer membrane ⍺-helical proteins play critical roles in mitochondrial-cytoplasmic communication, but the rules governing the targeting and insertion of these biophysically diverse substrates remain unknown. Here, we first defined the complement of required mammalian biogenesis machinery through genome-wide CRISPRi screens using topologically distinct membrane proteins. Systematic analysis of nine identified factors across 21 diverse ⍺-helical substrates reveals that these components are organized into distinct targeting pathways which act on substrates based on their topology. NAC is required for efficient targeting of polytopic proteins whereas signal-anchored proteins require TTC1, a novel cytosolic chaperone which physically engages substrates. Biochemical and mutational studies reveal that TTC1 employs a conserved TPR domain and a hydrophobic groove in its C-terminal domain to support substrate solubilization and insertion into mitochondria. Thus, targeting of diverse mitochondrial membrane proteins is achieved through topological triaging in the cytosol using principles with similarities to ER membrane protein biogenesis systems.

## INTRODUCTION

α-helical integral membrane proteins are found in prokaryotic plasma membranes as well as across eukaryotic membranes. In eukaryotes, this class of proteins is primarily found throughout the secretory pathway and in mitochondrial membranes where they play critical roles in maintaining cellular homeostasis and communication (von Heijne, G., 2007, Rapaport et. al, 2017, Hegde and Keenan, 2022). Their diversity in function is enabled by a wide range in physical properties such as topology, and key features of transmembrane domains (TMDs) such as hydrophobicity, charged residues, and variable helix lengths. This biophysical diversity poses particular challenges for transmembrane protein biosynthesis and folding (Krogh et. al, 2001, Wickner and Schekman, 2005, Rapoport, 2007). Additionally, the presence of aggregation prone hydrophobic TMDs means eukaryotic cells must develop pathways that can both keep these proteins soluble in the cytosol and deliver them to the correct compartment (White and von Heijne, 2008). Ultimately, the cytosolic biosynthetic machinery must coordinate with membrane embedded machinery to facilitate the insertion and folding of α-helical membrane proteins in the lipid bilayer.

Eukaryotic membrane protein biogenesis has been most intensely studied at the endoplasmic reticulum (ER), which is the site of biogenesis for most membrane proteins (Shao and Hegde, 2011). In order to accommodate the diversity of proteins translocated into the ER, the cell has developed parallel pathways that cater to different substrate types. Initially, it was thought that these proteins all relied on co-translational substrate targeting mediated by the ribosome-associated signal recognition particle (SRP) which shields nascent TMDs (Keenan et. al, 2001, Halic et. al, 2004, Voorhees and Hegde, 2015) while delivering them to the SEC61 translocon for insertion into the membrane (Heinrich et. al, 2000, Weng et. al, 2021). However, it is now established that SRP and SEC61 cannot accommodate the full diversity of ER membrane proteins. For example, tail-anchored proteins, defined by a single TMD close to the stop codon, are necessarily targeted post-translationally as the nascent chain is released prior to the emergence of the TMD from the ribosomal tunnel (Nyathi et. al, 2013, Zhang and Shan, 2014, Guna et. al, 2023). After release from the ribosome, cytosolic factors such as GET3, via SGTA (Wang et. al, 2010) and calmodulin (Guna et. al, 2018), capture TAs based on hydrophobicity, and subsequently target them to either the EMC or GET complexes for insertion at the ER (Wang et. al, 2014, Guna et. al, 2018, Chitwood et. al, 2018, McDowell et. al, 2020). ER membrane protein biogenesis is thus supported by a complex network of factors with distinct targeting and insertion pathways required to handle specific types of substrates, also utilizing TMD assembly factors (Chitwood and Hegde, 2020), and quality control factors (Rodrigo-Brenni et. al, 2014, Shao et. al, 2017, McKenna et. al, 2020) to degrade aberrant TMDs. Whether a similar set of strategies exists for α-helical protein biogenesis at the mitochondria, a less well-studied system, remains unknown.

The outer mitochondrial membrane (OMM) proteome is critical for mitochondrial-cytoplasmic communication. There are two central classes of integral OMM proteins – β-barrel proteins, which are also found in bacterial outer membranes, and α-helical TMD containing proteins, an evolutionarily newer and broader class. The OMM α-helical proteins resemble the ER integral membrane proteins in their biophysical and topological diversity, and they can be broadly categorized by type as signal-anchored (SA, single TMD anchored at the N-terminus), tail-anchored (TA, single TMD anchored at the C-terminus), and topologically distinct polytopic proteins. This similarity suggests the existence of multiple distinct α-helical targeting and insertion pathways at the OMM, but mechanistic knowledge of the biogenesis process, including the full range of factors involved, remains incomplete.

All OMM proteins are nuclear-encoded and synthesized in the cytosol and must be specifically targeted to the membrane in an insertion competent manner (Becker, Song, and Pfanner, 2019, Gupta and Becker, 2021). Unlike the majority of mitochondrial proteins that are destined for the intermembrane space (IMS), inner membrane or matrix, OMM proteins do not have canonical mitochondrial targeting sequences (MTS). Elegant studies have defined the import pathways for MTS proteins, which are imported into the mitochondria through the central TOM40 pore (Hill et. al, 1998) in yeast and mammals. First, they are recognized by the TOM20 and TOM22 receptors (Abe et. al, 2000, Su et. al, 2022). Hydrophobic internal carrier proteins are recognized by the alternate TOM70 receptor (Backes et. al, 2018) before import through TOM40. β-barrel OMM proteins are also initially imported through TOM40 and subsequently inserted from the IMS into the membrane by the SAM complex using a mechanism evolutionarily conserved from bacteria (Kutik et. al, 2008, Takeda et. al, 2021).

By contrast, α-helical outer membrane proteins do not use either the TOM40 pore or the SAM complex (Becker et. al, 2011, Doan et. al, 2020) and structural studies have established that, unlike the ER insertases, TOM40 does not provide an energetically favorable lateral gate for membrane protein insertion (Bausewein et. al, 2017, Tucker and Park, 2019, Araiso et. al, 2019, Wang et. al, 2020). Instead, recent studies have established novel insertases for α-helical outer membrane proteins in yeast, trypanosomes and mammals (Becker et. al, 2011, Vitali et. al, 2018, Guna et. al, 2022). The MIM complex was identified as an α-helical insertase in yeast (Becker et. al, 2011, Krüger et. al, 2017, Doan et. al, 2020) and later shown to be functionally interchangeable with the trypanosome multi-spanning protein pATOM36 (Vital et. al, 2018) despite a lack of structural or sequence similarity in an intriguing display of convergent evolution. More recently, based on genetic, structural and biochemical studies in human cells, we identified the multi-spanning α-helical proteins MTCH1/2 (Guna et. al, 2022) as being necessary and sufficient for inserting α-helical proteins in the outer mitochondrial membrane. However, MTCH2 has minimal exposure to the cytosol, and α-helical OMM proteins have varying dependencies on these insertases. Thus, there remain open questions regarding the nature of the cytosolic machinery responsible for targeting α-helical proteins to the OMM, and the rules used by this machinery to support the biogenesis of the full range of OMM proteins.

Here we employ a combination of systematic large-scale genetic screens and biochemical characterization of biogenesis factors to decipher the molecular logic involved in coordinating diverse α-helical substrate biogenesis at the mitochondrial outer membrane. First, we use genome-wide FACS-based CRISPRi screens in human cells for four topologically distinct outer membrane α-helical proteins to uncover a wide repertoire of mammalian biogenesis and quality control machinery. Then, we systematically test the effects of nine key identified factors across a diverse suite of twenty-one α-helical proteins in a targeted arrayed CRISPRi screen to determine how distinct biogenesis factors impact specific classes of helical OMM substrates. Finally, we characterize novel modes of cytosolic targeting to the outer membrane using biochemical and mutational studies based on high-confidence alpha-fold models and cell biological assays, showing that distinct chaperone networks triage α-helical proteins based on their topologies.

## RESULTS

### Systematic genome-wide CRISPRi screens identify factors required for α-helical OMM protein biogenesis

To uncover the machinery involved in diverse α-helical OMM protein biogenesis, we selected representative substrates of varying topologies to use as reporters in genome-wide CRISPRi screens: the iron-sulfur cluster biogenesis factor CISD1 as a SA protein (Wiley et. al, 2007, Paddock et. al, 2007, Wang et. al, 2017), the 3-TMD containing mitophagy effector FUNDC1 with its C-terminus in the IMS and N-terminus in the cytosol (Liu et. al, 2012), and the 5-TMD containing cholesterol translocator TSPO with its N-terminus in the IMS and C-terminus in the cytosol (Delavoie et. al, 2003, Liu et. al, 2006) (Figure 1A). As a basis for comparison, we used our recently described TA protein OMP25 (Guna et. al, 2022) which is known to be post-translationally targeted to the membrane and depends on MTCH2 for its insertion into the mitochondrial outer membrane. To systematically identify factors required for the biogenesis of CISD1, FUNDC1 and TSPO, we adapted a split-GFP reporter system (Vasseur et. al, 2021, Guna et. al, 2022) to a FACS-based screening platform. For this, we expressed GFP1-10 in the mitochondrial IMS and fused GFP11 onto the reporter terminus with predicted localization in the IMS (Figure 1A). This strategy allowed us to specifically monitor substrate integration into the outer membrane, rather than mis-localization into other organelles or retention in the cytosol. Therefore, topologically correct insertion of our substrates into the OMM should result in complementation and GFP fluorescence. Using microscopy, we verified mitochondrial localization of our reporters (Figure S1A) and confirmed topologically accurate insertion by showing that only the reporters fused to GFP11 at the terminus predicted to be in the IMS result in fluorescence (Figure S1B).

**Figure 1.**
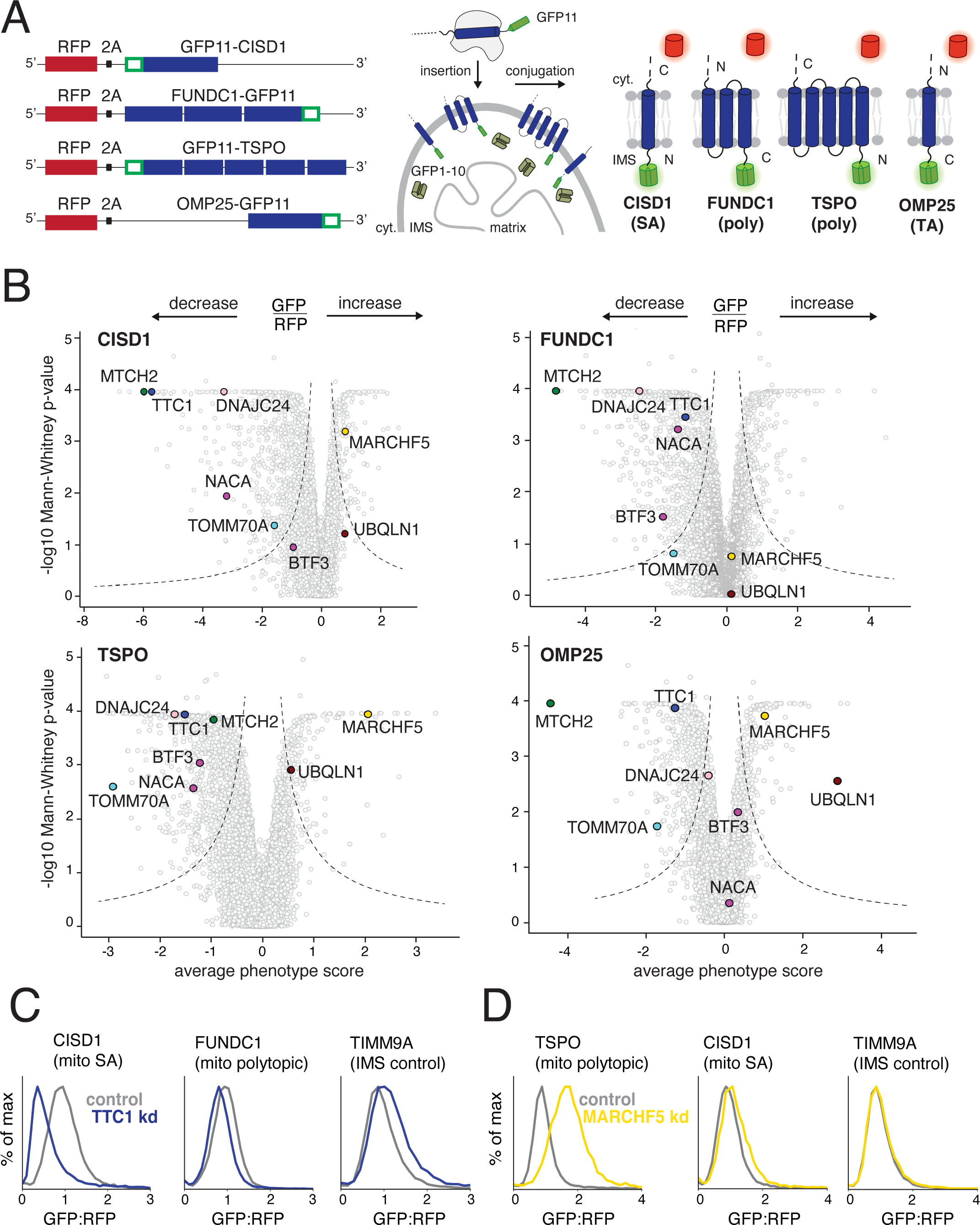
Identifying factors required for the biogenesis of topologically distinct α-helical mitochondrial outer membrane proteins. (A) The insertion of α-helical mitochondrial outer membrane proteins of diverse topologies (SA: signal-anchored, poly: polytopic and TA: tail-anchored) was queried using a split GFP reporter system. GFP11-fused reporters were expressed in a K562 cell line constitutively expressing GFP1-10 in the intermembrane space (IMS) such that insertion in the correct topology would result in complementation and GFP fluorescence. RFP (red fluorescent protein) separated by a viral P2A acts as a translation normalization marker. (B) Volcano plots of the GFP:RFP stabilization phenotype for the three strongest sgRNAs and Mann-Whitney p-values for the genome-wide CRISPRi FACS screens (two independent replicates each) for the four reporters in (A). Each gene is indicated by a gray dot, with specific genes of interest that either decrease or increase the GFP:RFP ratio highlighted across all four screens. The OMP25 screen was conducted in a previous study with identical screening conditions and is replotted to allow for direct comparison (Guna et. al, 2022). (C) Integration of the indicated GFP11-fused reporters in K562 cell lines expressing IMS GFP1-10 in the presence of a non-targeting (control) sgRNA or sgRNAs targeting TTC1 or (D) MARCHF5. TIMM9A is an IMS control reporter. Flow data was normalized based on the GFP:RFP ratio and plotted as histograms. See Figures S1 and S2.

We then engineered monoclonal K562 cell lines to stably express the full-length endogenous sequences of each of our reporters along with an expression normalization marker (RFP), as well as IMS localized GFP1-10 and the CRISPR inhibition machinery (Figure S1C) to enable genome-wide pooled screening. After transduction with a genome-scale CRISPRi sgRNA library into individual reporter cell lines, (Horlbeck et. al, 2016), cells with perturbed GFP/RFP fluorescence were isolated using FACS and their associated sgRNAs were identified by next-generation sequencing. We reasoned that by using this ratiometric approach which has previously been successful in identifying OMM α-helical biogenesis factors (Guna et. al, 2022), we could analyze sgRNAs enriched in either high or low populations of GFP/RFP fluorescence to identify putative quality control and biogenesis factors, respectively.

We first identified several factors previously known to impact biogenesis of our reporters as “hit” genes, thus validating our screening approach. Primarily, MTCH2 depletion clearly impacted all our substrates albeit to varying degrees, confirming its ability promote the insertion of biophysically diverse α-helical proteins beyond TAs (Guna et. al, 2022) (Figure 1B). Additionally, knockdown of the alternate TOM complex receptor TOMM70 (gene name TOMM70A) specifically strongly decreased biogenesis of the polytopic TSPO reporter, consistent with its role in mediating TSPO biogenesis (Otera et. al, 2007) (Figure 1B). Further support of our screen design came from CISD1 where depletion of its partner iron sulfur-biogenesis factors (FAM96A, NARFL, NFS1, ABCB7) specifically impacted CISD1 OMM stability (Figure S2A; Dubreuil et. al, 2020).

Importantly, we also identified putative biogenesis and quality control factors not previously associated with α-helical OMM proteins, and many of these factors showed strong differential impacts on the four substrates explored here. Cytosolic factors were prominent among these, presumably mediating chaperoning and targeting to the outer membrane. Key among the cytosolic machinery was the poorly characterized TPR-domain containing protein TTC1, implicated potentially in protein folding and trafficking (Lotz et. al, 2008, Coukos et. al, 2021), but not specifically in mitochondrial outer membrane protein biogenesis. Depletion of TTC1 specifically impacted biogenesis of our SA reporter CISD1 (Figure 1B) without impacting the insertion of the topologically different outer membrane reporters or insertion into the IMS (Figure 1C). Given the ability of TTC1’s TPR domain to recruit cytosolic HSP90 or HSP70 (Schuefler et. al, 2000), it is an attractive candidate for chaperoning nascent substrates. Similarly, depletion of J-domain HSP40 chaperone DNAJC24 (Thakur et. al, 2012) impacted polytopic and SA reporters, but not the TA reporter. Intriguingly, both members of the co-translational chaperone NAC (NACA and BTF3) (Wiedmann et. al, 1994) were required for efficient biogenesis of polytopic reporters TSPO and FUNDC1, but their loss had no effect on TA reporter OMP25 (Figure 1B), suggestive of a need for these highly hydrophobic substrates to be shielded soon after emergence from the ribosome as is the case for ER polytopic proteins (Weng et. al, 2021).

We also identified factors whose depletion leads to increased reporter integration into the OMM, indicating possible roles in quality control. For example, depletion of membrane-resident E3 ligase MARCHF5 (Yonashiro et. al, 2006, Nagashima et. al, 2014) specifically impacted the outer membrane levels of the polytopic reporter TSPO (Figure 1B) but not SA screen reporter CISD1 or IMS control reporter TIMM9A (Figure 1D). Presumably, MARCHF5 ubiquitinates misfolded or aberrant TMDs of TSPO for proteasomal degradation, as shown for its known outer membrane substrates (Yonashiro et. al, 2006, Nagashima et. al, 2014). Additionally, we confirmed that UBQLN1 depletion specifically impacted the TA reporter OMP25 (Figure 1B, Guna et. al, 2022), but not those with other topologies, consistent with its role in facilitating degradation of tail-anchored mitochondrial membrane proteins that fail to be inserted (Itakura et. al, 2016). Comparing the phenotype scores of hits across the four different reporters screened showed differential effects (Figure S2A), supporting our initial hypothesis that a complex network of factors would likely be required to coordinate the diverse suite of outer membrane proteins. With this method of analysis, we also noted that canonical TOM complex receptors TOMM20 and TOMM22 (Su et. al, 2022) impacted our reporters to similar extents and tied this to general effects on the GFP1-10 system in the IMS (Figures S2B and S2C). Thus, this comparison also allowed us to rule out factors that simply modulate the IMS GFP1-10 system and are thereby not of further interest in relation to outer membrane protein biogenesis.

### Substrate type dictates factor dependence

We next sought to determine whether factors identified in our genome-wide CRISPRi screens had reporter-specific effects, or more general roles in α-helical OMM protein biogenesis. Our limited reporter set for the screens made it difficult to define mechanistic patterns and necessitated a more comprehensive approach. To delineate the logic by which our hits impact diverse α-helical OMM proteins, we designed a comprehensive panel of endogenous α-helical OMM reporters fused to GFP11 in the RFP-P2A reporter cassette (Figure 1A). These included functionally diverse SA, TA and polytopic proteins. Within polytopic proteins, we also selected for those varying in number of TMDs and orientations of the termini within the membrane (Figures 2A and S3A). For certain proteins, such as MUL1 and MARCHF5, which were not compatible with the split GFP system due to having both termini in the cytosol, we fused the endogenous sequences to full-length GFP at the C termini and validated correct localization by microscopy. As controls we used IMS-localized proteins that either use canonical targeting sequences, such as MICU1 and LACTB, or have internal targeting sequences and are dependent on known machinery, such as TIMM9A and TIMM10 which are dependent on MIA40 (Mesecke et. al, 2005, Milenkovic et. al, 2009).

**Figure 2.**
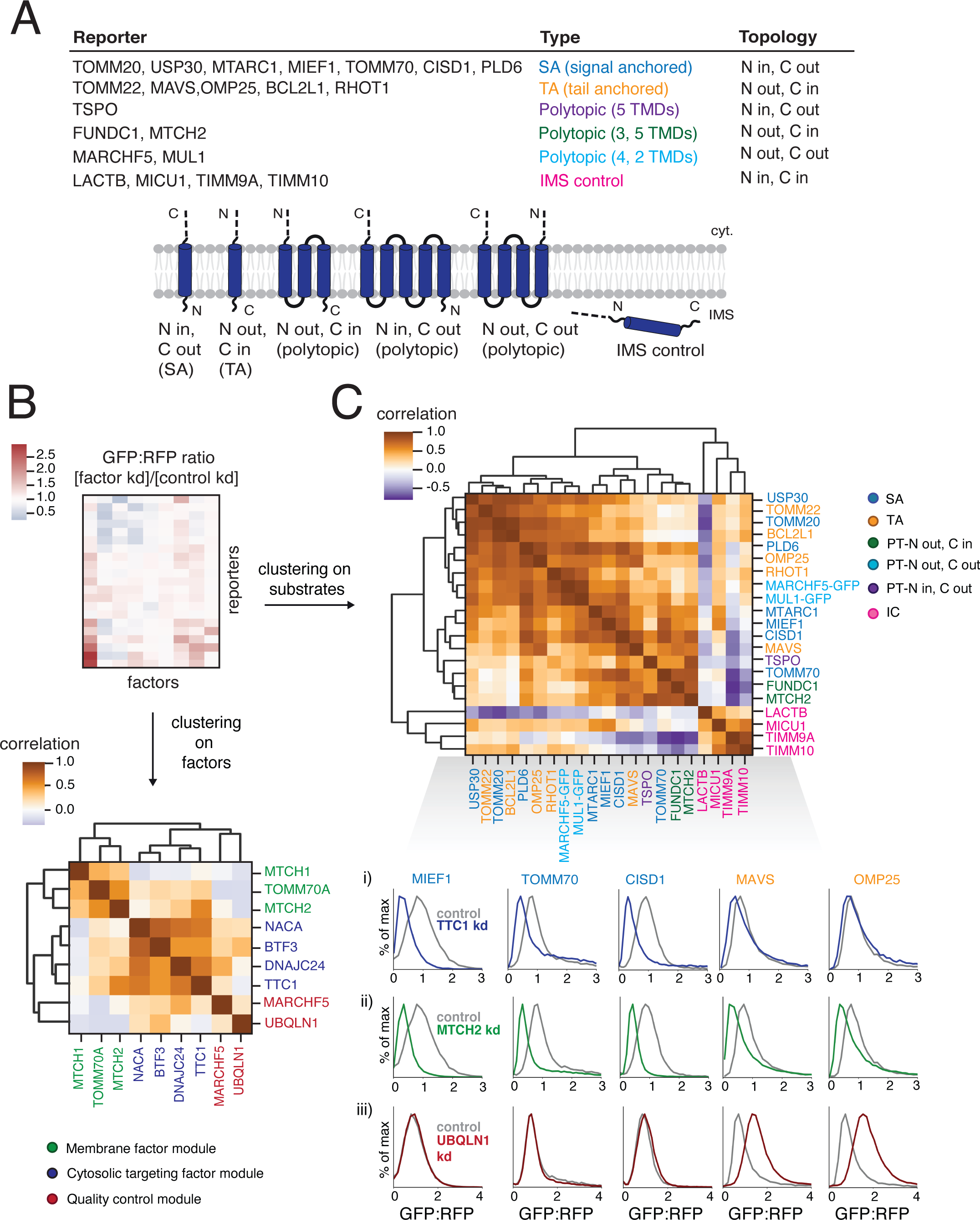
Systematically exploring pathways affecting the biogenesis of diverse α-helical mitochondrial outer membrane proteins. (A) Table describing a comprehensive set of α-helical outer membrane proteins used as reporters to query the specificity of hits (highlighted in Figure 1B) derived from the genome-wide CRISPRi screens. Type and topology of each protein is stated and depicted. IMS control reporters were also included (for a total of 21 substrates assayed). ‘LACTB’ and ‘MICU1’ indicate just the respective targeting sequence fused to GFP11. For proteins with both termini in the cytosol (MUL1 and MARCHF5) and therefore incompatible with the split GFP system, full-length GFP was fused to the C-termini. (B) To analyze the effects of factors of interest on α-helical proteins of different topologies, each reporter in (A) was expressed in K562 ZIM3 CRISPRi cells constitutively expressing IMS GFP1-10 and a sgRNA guide against the factor or a non-targeting control and analyzed by flow cytometry. The GFP:RFP ratios were then normalized to the non-targeting control ratios to generate a comprehensive dataset represented as a heatmap (see Figure S3A for details). Hierarchical clustering of the correlation matrix of all factors results in the assignment of putative biogenesis pathways. (C) Clustering of the correlation matrix of all reporters assigns outer membrane reporters into clusters largely defined by type and topology. Individual histograms showing GFP:RFP changes in response to factor depletion that are predictive of SA and TA reporter clustering patterns are shown here: i) responses to TTC1 depletion, ii) responses to MTCH2 depletion, iii) responses to UBQLN1 depletion. See Figure S3 for complete reporter GFP:RFP changes (for all factors). See Figures S3 and S4.

Due to their differential phenotypic effects in the screens, we chose to test the cytosolic factors TTC1, NACA, BTF3 and DNAJC24 and the putative quality control factors MARCHF5 and UBQLN1 across the entire reporter panel. Additionally, we included insertases MTCH1 and MTCH2 and alternate TOM complex receptor TOMM70, which also appear to differentially direct outer membrane integration of biophysically diverse α-helical proteins. We then conducted a large arrayed screen in K562 CRISPRi cells constitutively expressing GFP1-10 in the IMS in duplicate. Each factor of interest was individually depleted and the expression of each reporter was assessed. These data yielded a 42 x 9 matrix (21 substrates tested in duplicate against 9 factor depletions) which we depicted as a heatmap colored by the fold change in reporter biogenesis (GFP:RFP ratio) for each factor perturbation relative to the control condition (non-targeting sgRNA) (Figure S3A, see also Figures 2C and S3B for representative flow cytometry plots from which the individual entries in this heat map are derived). Depletion of each factor was confirmed using qRT-PCR (Figure S3C). Notably, each factor had differential effects across the reporter set, demonstrating a lack of general dysregulation of outer membrane protein biogenesis. These effects were also not driven by an impact on the GFP1-10 system as the biogenesis of the IMS controls was relatively unchanged across all factor perturbations (heatmap in Figure S3A; table S5). We only observed minor biogenesis defects in MICU1 and LACTB-targeted GFP11 reporters under DNAJC24 depletion (Figure S3A; table S5).

Unbiased hierarchical clustering on the heatmap raw data (Figure S3A) revealed distinct patterns of biological interest. First, the reporters were largely organized by type (i.e., SA, TA, polytopic and IMS control) as well as topology within type in the polytopic proteins – for example, FUNDC1 and MTCH2 share the same topology and cluster closest to each other away from TSPO. Our analysis largely partitioned the signal-anchored (SA) proteins by their differential dependencies on TTC1, as SA reporters showed the strongest biogenesis defects upon TTC1 depletion with an ∼50% reduction in OMM integration (Figure S3A). The two components of the NAC complex, NACA and BTF3, primarily impacted the polytopic reporters TSPO, FUNDC1 and MTCH2, helping distinguish them from other substrate types. We also noted phenotypic effects of several factors that did not appear to solely correspond to substrate type. Among the quality control factors, MARCHF5 depletion specifically impacted TSPO and other single-pass reporters such as RHOT1 (TA) and PLD6 (SA), but not other polytopic reporters FUNDC1 and MTCH2. Similarly, TOMM70 depletion reduced TSPO OMM integration by ∼70% but no other reporter to the same extent – this combined with MARCHF5’s unique effect on TSPO separated it from other polytopic reporters FUNDC1 and MTCH2 (Figures S3A and S3B). Finally, we saw that NACA separated from the rest of the factors in the heatmap, likely due to previously observed toxicity in its absence (Hotokezaka et. al, 2009) that is amplified with ZIM3-mediated CRISPRi depletion (Alerasool et. al, 2020; Figures S3A and S3C). The toxicity caused a wider distribution of the GFP signal peak due to a much smaller number of recorded live cell events which inflated the GFP:RFP ratios across the reporter set and led to inaccurate measurements of reporter biogenesis. To resolve this, we used KOX1-mediated weaker CRISPRi depletion of NACA (see Figures 4A and S6A), which replicated the BTF3 depletion results in primarily impacting polytopic reporters.

**Figure 3.**
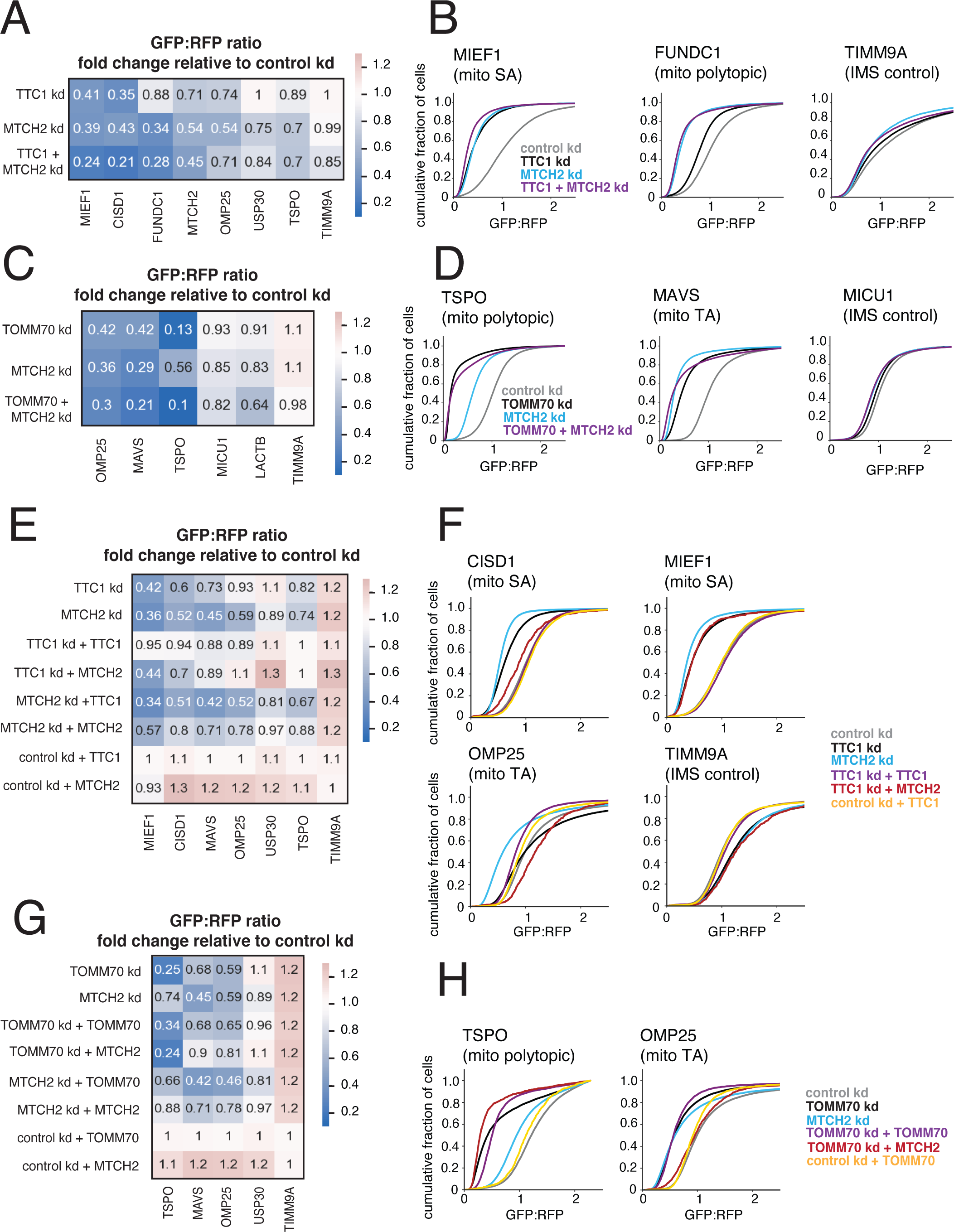
TTC1 and TOMM70 exert effects on α-helical membrane protein integration independent of their effects on the MTCH2 insertase. (A) Arrayed dual guides were used to assess the genetic interaction of TTC1 and MTCH2 by monitoring outer membrane integration of GFP11-fused α-helical reporters dependent on TTC1 and MTCH2 (MIEF1, CISD1, MTCH2), MTCH2 alone (USP30, FUNDC1, OMP25) and neither (TSPO, TIMM9A) (dependencies noted from Figure S3A). Reporters were expressed in K562 ZIM3 CRISPRi IMS GFP1-10 cells and i) sgRNAs targeting TTC1 and MTCH2, ii) sgRNAs targeting TTC1 and a non-targeting control, iii) sgRNAs targeting MTCH2 and a non-targeting control. Data is represented as a heatmap colored by the fold change in reporter integration (GFP:RFP ratio) for each condition relative to cells expressing a non-targeting sgRNA. (B) Flow data for representative reporters from (A) plotted as cdf (cumulative distribution function) plots. (C) The genetic interaction of TOMM70A and MTCH2 was assessed in a similar manner to (A) with reporters dependent on TOMM70A and MTCH2 (OMP25, MAVS), TOMM70A alone (TSPO) and neither (MICU1, LACTB, TIMM9A). As in (A), dependencies are noted from arrayed screen results in Figure S3A. (D) Flow data for representative reporters from (C) plotted as cdf plots. (E) To determine the extent to which TTC1 biogenesis defects are a secondary consequence of MTCH2 loss, outer membrane integration of GFP11-fused α-helical reporters was measured in K562 IMS GFP1-10 cells constitutively expressing a sgRNA against TTC1 and i) exogenous 3xFLAG-tagged TTC1 in a cassette with a BFP translation marker, ii) exogenous MTCH2 with a BFP translation marker and iii) BFP alone. Reporter integration was measured with the same conditions for cells expressing sgRNAs targeting MTCH2 and non-targeting sgRNAs (control). Data is represented as a heatmap colored in the same manner as (A) and (C). (F) Flow data for representative reporters from (E) plotted as cdf plots. (G) The extent to which TOMM70A biogenesis defects are a secondary consequence of MTCH2 loss is assessed in the same manner as (E). (H) Flow data for representative reporters from (E) plotted as cdf plots. See Figure S5.

**Figure 4.**
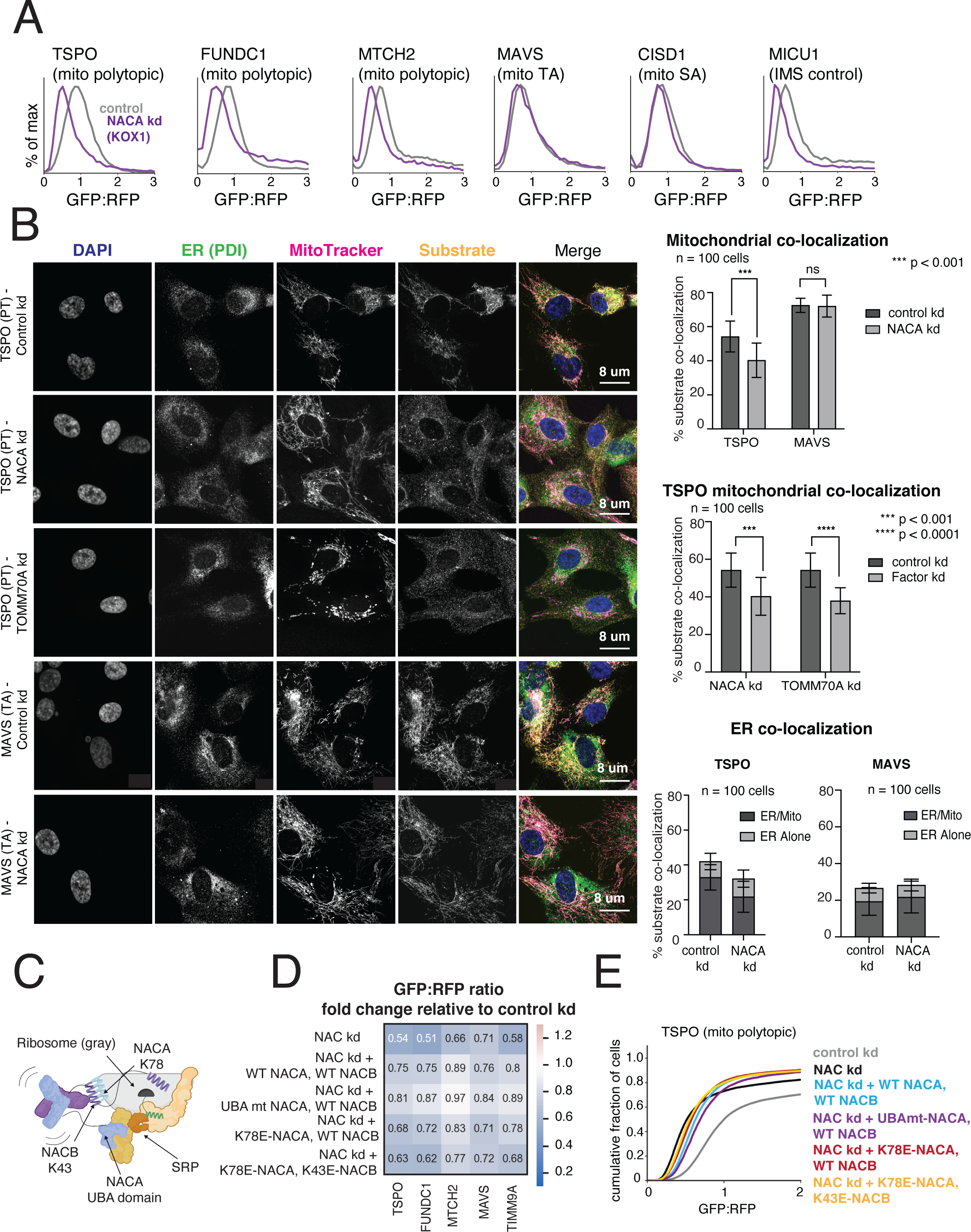
The NAC complex is required for the efficient delivery of polytopic α-helical proteins to the mitochondrial outer membrane. (A) Integration of indicated GFP11-fused SA, TA, polytopic and IMS control reporters in K562 KOX1 CRISPRi cells constitutively expressing IMS GFP1-10 and either a sgRNA targeting NACα or a non-targeting sgRNA (control). (B) (Left) Changes in localization of endogenous TSPO (polytopic) and MAVS (TA) were assessed using immunostaining and confocal microscopy in RPE1 ZIM3 CRISPRi cells expressing sgRNAs targeting NACα, TOMM70 or a non-targeting control. Merge colors: Blue: DAPI, Green: ER, Yellow: Substrate, Magenta: MitoTracker. (Right) Quantification of endogenous substrates co-localized to MitoTracker (mitochondrial marker) and PDI (ER marker) were measured and plotted (see Methods for details). Quantification of substrates co-localized to the ER were split into i) percent co-localized to ER/MitoTracker overlapping regions and ii) percent co-localized to the rest of the ER (‘ER alone’) (see FIgure S6D). For more detailed substrate localization analysis across all conditions, see Figures S6C-F. Error bars show mean ± SD of 100 cells. Statistical significance was evaluated by multiple unpaired t-tests with the Holm-Sidak multiple test correction. ***, p < 0.001. ****, p < 0.0001. ns (non-significant), p > 0.05. (C) Model of a translating ribosome engaging NAC and SRP. NACα (blue) and NACβ (purple) interact with the ribosome (gray) through electrostatic interactions mediated by the indicated lysine residues (NACα K78 and NACβ K43). SRP is recruited to the ribosome through electrostatic interactions with the UBA domain of NACα (at residues D205 and N208), where it engages the substrate nascent chain (green) co-translationally. (D) The effects of ribosome-binding mutants of NACα and NACβ (K78E and K43E respectively) and a UBA-domain binding mutant of NACα (D205R + N208R) on integration of select GFP11-fused reporters in K562 KOX1 CRISPRi IMS GFP1-10 cells was tested using a similar strategy as in Figures 3E and 3G. Both members of the NAC complex were simultaneously depleted using CRISPRi and abilities of exogenous wild-type (WT) and mutant versions of NACα and NACβ to rescue reporter integration defects were assessed. Rescue constructs are in a cassette with a BFP translation marker. Data is represented as a heatmap colored by the effects of exogenous additions of NACα and NACβ (WT and mutant versions as indicated) on reporter integration in the mitochondria relative to the control condition (non-targeting sgRNA + BFP alone, as shown in Figures 3E and 3G). (E) Flow cytometry data for polytopic reporter TSPO across all conditions shown in (D) represented as a cdf plot. See Figures S6 and S7.

We next sought to further organize our factors of interest into putative pathways by constructing a correlation matrix and subsequently performing hierarchical clustering (Figure 2B). This allowed us to derive patterns of factor utilization and revealed putative biogenesis ‘modules’ of factors that function together. The individual correlation values were highly reproducible across replicates (Figure S4A) and the resulting matrix can be clustered into three broad putative categories: i) a quality control module comprising MARCHF5 and UBQLN1, ii) a cytosolic targeting module comprising TTC1, DNAJC24, NACA and BTF3 and iii) a membrane factor module comprising MTCH1, MTCH2 and TOMM70 (Figure 2B). Within the cytosolic targeting module, both members of the co-translational NAC complex NACA and BTF3 clustered closest to each other, segregating from the other targeting factors TTC1 and DNAJC24, which have both been implicated as having HSP co-chaperoning function (Schuefler et. al, 2000, Thakur et. al, 2012) (Figure 2B). Looking at individual correlation values, we found that the closest correlating factor pair in the entire dataset was NACA and BTF3 (Figures 2B and S4A). This validated our clustering analysis as both are known to work together in the same protein complex in co-translational chaperoning and targeting (Shen et. al, 2019, Jomaa et. al, 2022). Aside from seeing that the factors cluster into distinct functional modules, we also noted interesting individual strong correlations, such as that of TTC1 and MTCH2 (Figure 2B). This likely results from their impacts on a largely similar set of reporters (Figure S3A) and raises the intriguing possibility that substrates targeted by TTC1 are recruited to MTCH2 for subsequent insertion.

Using the same strategy, we clustered all the reporters by their patterns of correlation with each other (Figure 2C) and showed that the correlation matrix is highly reproducible (Figure S4B). Once again, we saw that the reporters were largely organized by type, and topology within type as described for the raw GFP:RFP fold change cluster map (Figure S3A). Importantly, we showed that some of the IMS controls, particularly LACTB, anti-correlate with the outer membrane reporters, reinforcing the uniqueness of the outer membrane protein biogenesis system (Figure 2C). As seen in the heatmap, separation of the SA and TA proteins is likely governed by differential dependencies on several biogenesis and quality control factors, highlighted in Figure 2C (see sub-panels i, ii and iii). We further saw patterns of ‘sub-clustering’ within reporter types – for example, we attributed the tight clustering of a subset of SA proteins MIEF1, CISD1, MTARC1 and TOMM70 to their shared strong dependencies on MTCH2 and TTC1 (Figures 2C, S3A), while TA proteins like OMP25, BCL2L1 and TOMM22 depend on MTCH2 but not TTC1. Notably, we observed that TA protein MAVS interrupts the SA cluster in our map, clustering away from the other TA reporters. This is likely due to its weak TTC1 dependency (Figure 2C panel i) (Figure S3A). However, it also correlates highly with the TA protein OMP25 as it shares the characteristics of being much more dependent on MTCH2 (Figure 2C panel ii) than TTC1 for biogenesis and being stabilized by depletion of cytosolic quality control factor UBQLN1 (Figure 2C panel iii). Thus, we showed that analyzing the correlation matrix of all the reporters provided two layers of information – first, the hierarchical clustering which largely separates the reporters by type and topology, and second, the individual correlation values across all pairs of reporters which can help explain some of the idiosyncrasies in reporter positioning within the cluster map order.

Overall, we systematically established multiple biogenesis modules for α-helical outer membrane proteins and began to uncover the logic by which these modules are employed. Substrate type and topology emerged as critical deterministic factors for triaging nascent outer membrane proteins during biogenesis analogous to the biogenesis model for the ER integral membrane proteome, with cytosolic targeting and quality control factors driving most of the specificity in substrate selection.

### The cytosolic factors are the primary drivers of biogenesis pathway architecture

While we saw differential effects of factor depletion on α-helical protein biogenesis by type in the genome-wide CRISPRi screens and arrayed screen, we wanted to ensure that this was not driven by ancillary effects. Specifically, we saw strong correlations of i) TTC1 and MTCH2 and ii) TOMM70 and MTCH2 in the factor correlation matrix (Figure 2B), which arose from overlap in the reporters affected (Figure S3A). A potentially confounding factor is the biogenesis defect of the MTCH2 reporter upon depletion of TTC1 and TOMM70 (Figure S3A), which raises the possibility that some of the downstream phenotypic effects of TTC1 and TOMM70 knockdown are driven by loss of the MTCH2 insertase. We hypothesized that some of the weaker biogenesis defects for TA reporters by TTC1 and TOMM70 depletion could be indirectly driven by a loss of MTCH2. To investigate this, we first looked at the epistatic relationship between TTC1 and MTCH2. We expressed constructs with sgRNAs targeting i) TTC1 alone, ii) MTCH2 alone, iii) TTC1 and MTCH2 in K562 ZIM3 CRISPRi IMS GFP1-10 cells (Figure S5A) and measured the resulting outer membrane integration of a subset of our reporters – these included reporters that are strongly dependent on both TTC1 and MTCH2 (CISD1, MIEF1, MTCH2), mostly dependent on MTCH2 (USP30, FUNDC1, OMP25) and dependent on neither (TSPO, TIMM9A) (Figure S3A). Depletion of targeted factors was confirmed through qRT-PCR and immunoblotting (Figure S5A). We showed that for reporters mostly dependent on MTCH2, there was little to no additive effect of depleting TTC1. However, for reporters dependent on both, we saw a clearly additive effect of TTC1 depletion (Figures 3A and 3B), implying a direct dependency on TTC1 for biogenesis. We then investigated the genetic interaction of TOMM70 and MTCH2 using the same approach (Figure S5B). For a reporter primarily dependent on TOMM70 such as TSPO, there was no additive effect of depleting MTCH2, indicating that TOMM70 likely has a direct role in biogenesis. However, for reporters that show mutual dependencies such as OMP25, MAVS and the LACTB-GFP11 reporter, we observed a synthetic biogenesis defect (Figures 3C and 3D). This supports the idea that for these substrates, there is some biogenesis role for both factors with MTCH2 being the primary driver, in agreement with the genome-wide CRISPRi screen results for OMP25 (Figure 1B).

A direct role for TTC1 in SA biogenesis beyond regulating MTCH2 levels was further supported by the observation that expression of exogenous MTCH2 was sufficient to rescue the biogenesis defect caused by TTC1 depletion for weakly dependent TA reporters, but not for strongly dependent SA reporters CISD1 and MIEF1 (Figures 3E and 3F) – which were only rescued upon expression of exogenous TTC1. Similarly, we observed that expression of exogenous MTCH2 was not sufficient to rescue the biogenesis defect for primary TOMM70 substrate TSPO but was sufficient to rescue weakly TOMM70 dependent TA reporters (Figures 3G and 3H). We assessed expression of exogenous factors through immunoblotting (Figure S5C). Cumulatively, our epistatic analyses suggest that TTC1 and TOMM70 directly impact the biogenesis of select outer membrane protein types, such as SA and polytopic proteins respectively, with weaker effects on TA proteins driven by loss of the MTCH2 insertase. In conjunction with the analyses from Figure 2, these data provide a broader picture of the logic employed in driving biogenesis pathway architecture. Specifically, the cytosolic biogenesis and quality control factors drive much of the pathway distinctions among the α-helical OMM protein classes, all of which typically employ the insertase MTCH2 as a common entry point to the membrane. Indeed, the NAC complex distinguishes the polytopic proteins from the single-pass (Figures 2 and S3), with the SAs distinguished from the TAs within the single-pass proteins by their direct dependency on TTC1. TOMM70, while membrane-resident, is also known to act as a chaperone factor through its TPR domain recruiting cytosolic HSPs (Backes et. al, 2021) and further distinguishes polytopic and TA proteins.

Taken together, these results suggest that it is the specific TMD chaperoning requirements of diverse protein types that necessitates this complex biogenesis network. To expand on this, we next sought to define the roles of our putative chaperones NAC and TTC1, as they were the two targeting factors primarily driving cytosolic triaging within the overall biogenesis structure through their distinct substrate specificities.

### The NAC complex is required for efficient delivery of polytopic α-helical proteins to the OMM

The nascent polypeptide-associated complex (NAC) is a ribosome-associated chaperone complex composed of proteins NACα (NACA) and NACβ (BTF3), which are evolutionarily conserved and broadly essential in all eukaryotes except yeast (Bloss et. al, 2003, Markesich et. al, 2000). NAC sits at the opening of the ribosomal exit tunnel where it is poised to engage emerging nascent chains and ensure the fidelity of their targeting to the correct organelle. Classically, it was known to prevent promiscuous signal recognition particle (SRP) binding to the ribosome and subsequent mistargeting of mitochondrial proteins to the ER (Wiedmann et. al, 1994, Lauring et. al, 1995, Gamerdinger et. al, 2015), but has also recently been implicated in facilitating proper ER protein targeting by recruiting SRP for the correct substrates (Jomaa et. al, 2022) and in controlling co-translational N-terminal methionine excision for cytosolic proteins (Gamerdinger et. al, 2023). NAC also independently chaperones disease-associated polyglutamine (PolyQ) repeat proteins to prevent aggregation (Shen et. al, 2019). Currently, NAC has been shown to prevent the mis-localization of mitochondrial proteins to the ER, but only for proteins containing canonical mitochondrial targeting sequences (MTS) (Gamerdinger et. al, 2015, Jomaa et. al, 2022). Given that α-helical proteins in the OMM do not have MTS sequences, the observation that depletion of NAC affects their biogenesis potentially extends the role of NAC. We sought to determine whether NAC facilitates direct mitochondrial protein targeting for these substrates or mainly ensures fidelity by preventing their inaccurate targeting to the ER.

We found that both members of the NAC complex (NACA and BTF3) selectively affect integration of polytopic substrates in the outer membrane, with minimal impact on SA and TA substrates (Figures 4A and S6A). For these studies, we used KOX1-mediated CRISPRi to reduce the toxicity resulting from depletion of NACA using the more potent ZIM3 KRAB domain (Replogle et. al, 2022, Alerasool et. al, 2020) (noted in Figure S3A). As a control, we replicated previous observations that NAC affects the targeting of MTS containing reporters with MICU1 and LACTB targeting sequence-GFP11 reporters impacted by its depletion (Figures 4A and S6A). Additionally, we used confocal microscopy to specifically observe localization of a MICU1 targeting sequence-GFP reporter following NACA depletion (Figure S6B). We saw cytosolic retention of the MICU1-GFP reporter, with a clear decrease in mitochondrial localization. While our flow cytometry reporter data points to specific effects on polytopic substrates by NAC, the impact we saw on localization of the IMS GFP1-10 targeting sequence makes the reporter system imperfect for drawing definitive conclusions regarding direct targeting by NAC. Therefore, we sought to study changes in localization of endogenous outer membrane substrates with confocal microscopy following NAC depletion, which also allowed us to monitor changes in substrate localization across the entire cell.

We therefore depleted NACA in RPE1 cells and measured changes in localization of two endogenous α-helical outer membrane proteins, the polytopic protein TSPO and the TA protein MAVS, with immunofluorescence against the native proteins. To provide a point of comparison, we depleted TOMM70 which is known to specifically impact the mitochondrial integration of TSPO and lead to its accumulation in the cytosol (Otera et. al, 2007). Cells were stained with MitoTracker to mark mitochondria, protein disulfide isomerase (PDI) to mark the ER, and DAPI to mark nuclei to permit co-localization analyses. We observed that the mitochondrial localization of endogenous TSPO was significantly diminished under NACA depletion with no change in the mitochondrial localization of endogenous MAVS, replicating our screen and flow cytometry reporter assay results (Figure 4B). Notably, the mitochondrial localization of endogenous TSPO was decreased to a similar extent by TOMM70 and NACA depletion (Figures 4B, S6E and S6F). Both NACA and TOMM70 depletion resulted in a punctate form of TSPO which appeared to be largely retained in the cytosol (Figures 4B and S6C). To further address whether endogenous substrates were mistargeted to the ER to any appreciable degree, we quantified localization of TSPO and MAVS to the ER. In our analyses, we segregated the ER into regions that overlapped with the mitochondria and those that did not. We found that there was no significant change in MAVS or TSPO localization to the ER under NACA depletion (Figures 4B and S6D). Specifically, we noted a significant decrease in TSPO localization to the regions of the ER that overlap with the mitochondria, but no changes in the localization to the rest of the ER for MAVS or TSPO (Figures 4B, and S6D-F). Thus, endogenous TSPO was largely retained in the cytosol under NACA depletion, consistent with a primary biogenesis defect. This was phenocopied by TOMM70 depletion except for a minor decrease in TSPO colocalization to the region of the ER that did not overlap with the mitochondria (Figure S6E), which could be due to a stronger primary biogenesis defect (Figure S3A).

We next sought to test the role of NAC’s ribosome-binding function in polytopic protein biogenesis for potential co-translational targeting, as well as any contribution of the SRP-binding UBA domain of NACA, shown to be essential for accurately targeting ER proteins (Jomaa et. al, 2022). Given that the primary function of the UBA domain is to target proteins to the ER with high fidelity, we reasoned that it would not be critical for direct mitochondrial targeting. Accordingly, we made point mutations in the ribosome-binding globular domains of NACα and NACβ in conserved lysine residues critical for rRNA interaction (Figure 4C; Jomaa et. al, 2022), as well as in the UBA domain of NACA in conserved residues mediating electrostatic interactions with SRP (Figure 4C; Jomaa et. al, 2022). We then determined the abilities of these mutants to rescue split-GFP reporter biogenesis defects from depleting both components of the NAC complex, relative to rescue by adding back both components of wild-type NAC in the K562 KOX1 CRISPRi IMS GFP1-10 cell line. Indeed, the integrity of the UBA domain did not appear to be critical, while the ribosome binding function influenced efficiency of reporter integration (Figures 4D, 4E and S7A). Exogenous expression of all NAC rescue constructs was assessed through immunoblotting (Figure S7B). Depletion of NACA and BTF3 with KOX1 and ZIM3 CRISPRi for reporter assays shown (Figures 4A, 4D, S6A) were measured using qRT-PCR (Figure S7C).

Cumulatively, we point to a role for NAC as a targeting factor with a selective effect on outer membrane polytopic proteins, facilitating direct substrate targeting to the mitochondria rather than simply preventing ER mis-localization. The role of NAC ribosome association for these substrates also raises the intriguing possibility that they are co-translationally targeted, which has been implicated in yeast mitochondria for MTS proteins (Fünfschilling and Rospert, 1999, Yogev et. al, 2007) with the receptor OM14 proposed to bind NAC at the outer membrane in cooperation with Sam37 and Tom70 (Lesnik et. al, 2014, Ponce-Rojas et. al, 2017). Further studies could establish whether this is indeed the case for mammalian mitochondria and to determine the mechanism of NAC-mediated outer membrane targeting.

### TTC1 is a novel mitochondrial outer membrane protein biogenesis factor

In contrast to polytopic proteins, single-pass outer membrane proteins were not impacted by NAC depletion (Figures 2, S3A and 4A). Rather, we saw from both our genome-wide screens and arrayed analyses that biogenesis of most SA proteins is strongly impacted by depletion of the cytosolic factor TTC1. TTC1 has a conserved tetratricopeptide repeat (TPR) domain implicated in binding HSP90 with a potential role in protein trafficking (Lotz et. al, 2008, Coukos et. al, 2021), but is otherwise largely uncharacterized. We sought to understand TTC1’s role more generally, as it has not previously been associated with mitochondrial protein biogenesis defects. We also wanted to determine if TTC1’s mitochondrial effects are specific and not related to defects at any other organelle, such as the ER. First, we studied the transcriptional response resulting from TTC1 perturbation to understand the broader cellular changes associated in an unbiased manner. Minimal distortion embedding of Perturb-seq data (Replogle and Saunders et. al, 2022) for genes with depletion causing a strong transcriptional response in K562 cells showed TTC1 strongly correlating with key mitochondrial import and biogenesis factors such as members of the TOM, TIM, SAM and MICOS complexes, suggesting a potential role for TTC1 in mitochondrial import (Figure S8A). Additionally, plotting integrated stress response (ISR) and unfolded protein response (UPR) activation scores for all essential genes in the genome-scale Perturb-seq dataset in K562 cells (Replogle and Saunders et. al, 2022) showed that TTC1 perturbation strongly activates the ISR without activating the UPR, distinct from ER biogenesis factors such as SRP68 (Figure S8B). We also observed that TTC1 depletion causes pronounced mitochondrial network fragmentation in RPE1 cells (Figure 5A) without any notable effects on ER morphology (figure 5B). Within the mitochondria, the direct effects of TTC1 on nascent reporter biogenesis and integration were limited to the outer membrane proteins (Figure S8C). Furthermore, we showed that ER-localized split-GFP reporters of varied topologies were not destabilized by TTC1 depletion in a K562 ER GFP1-10 CRISPRi cell line (Figures S8D-E). We also showed a significant mitochondrial respiration defect caused by TTC1 depletion which was rescued by adding back exogenous TTC1 (Figure 5C). Importantly, we established that TTC1 impacts mitochondrial integrity directly rather than indirectly by affecting MTCH2 levels (Figures S8F-G), as MTCH2 levels are also known to regulate mitochondrial morphology and fragmentation (Labbé et. al, 2021).

**Figure 5.**
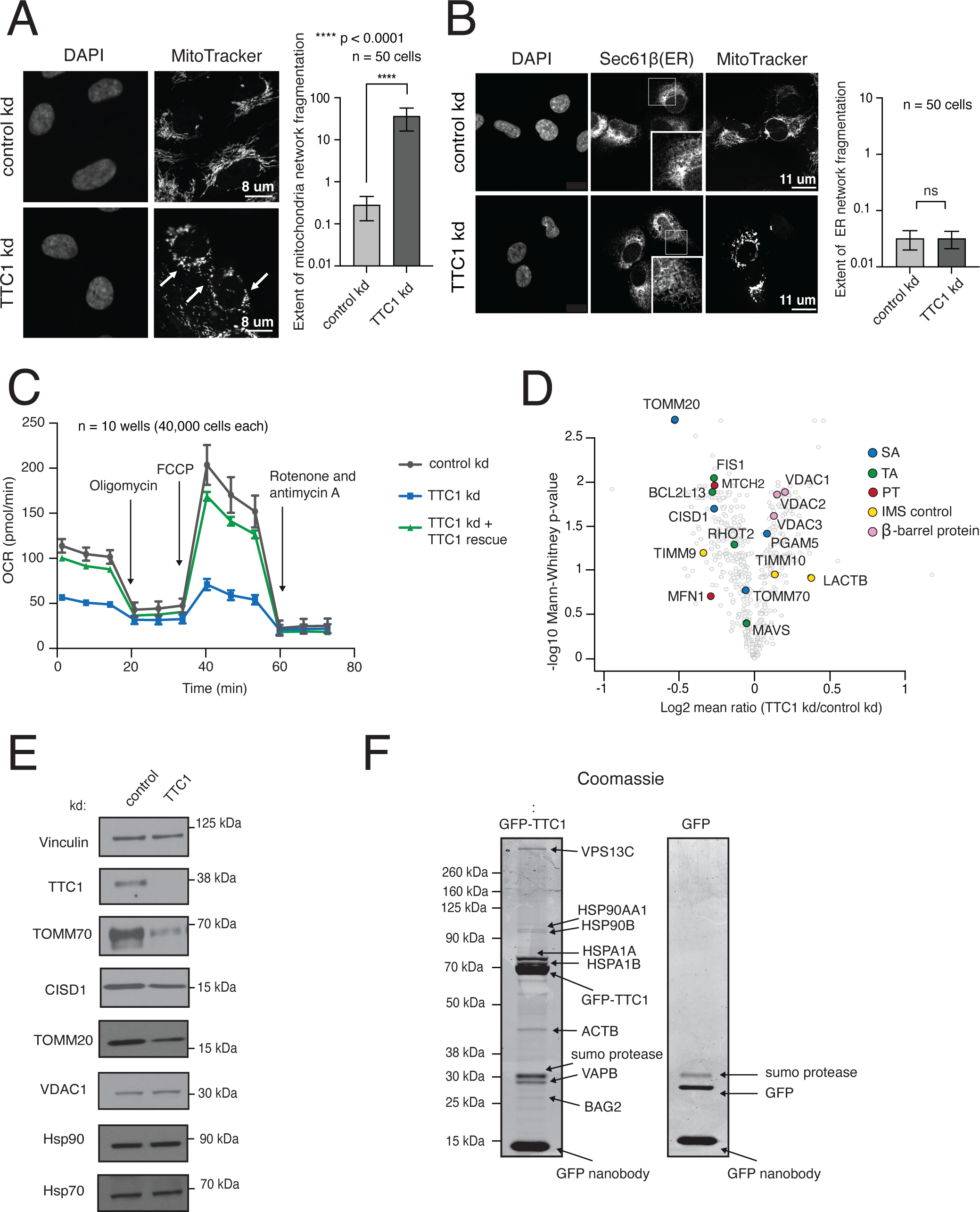
TTC1 is required for mitochondrial integrity and the biogenesis of endogenous outer membrane proteins. (A) RPE1 ZIM3 CRISPRi cells expressing either a sgRNA targeting TTC1 or a non-targeting sgRNA were analyzed by super-resolution confocal microscopy to assess changes in mitochondrial morphology. The white arrows indicate mitochondrial fragmentation in TTC1 depleted cells. Extent of network fragmentation is calculated using the median count: area ratio of the mitochondrial network in each condition with log10 values plotted. (B) The effects of TTC1 depletion on ER morphology in RPE1 ZIM3 CRISPRi cells, assessed through immunostaining with Sec61β and calculated as in (A). For (A) and (B), error bars show mean ± SD of 50 cells. Statistical significance was evaluated using the Welch corrected unpaired t-test. ****, p < 0.0001. ns (non-significant), p > 0.05. (C) Mitochondrial respiration was assessed in K562 ZIM3 CRISPRi cells depleted of TTC1 where levels were then rescued with an exogenous copy (see Figure S10D for protein levels) using a Seahorse assay to monitor oxygen consumption rates (OCR). Arrows indicate time of drug treatment. Data are presented as average ± SD, n = 10. (D) TMT mass-spectrometry analysis of K562 KOX1 CRISPRi cells expressing guides targeting TTC1 relative to cells expressing non-targeting guides (control). Data shown here are normalized to mitochondrial proteome levels and evaluated for statistical significance across 3 biological replicates. Outer membrane proteins of interest are colored by topology. (E) Immunoblotting of endogenous proteins in K562 KOX1 CRISPRi TTC1 depleted cells and control cells (non-targeting sgRNA) using vinculin as the loading control. (F) HEK293T cells stably expressing endogenously tagged TTC1-GFP were used to identify endogenous TTC1 interaction partners. TTC1-GFP was purified under native conditions using an anti-GFP nanobody (Pleiner et al, 2020). Interaction partners were analyzed by separation through SDS-PAGE, detection with Coomassie staining and identification by mass spectrometry. As a control for non-specific interactions, HEK293T cells were transduced with GFP before purification using the GFP-nanobody and downstream analysis in the same manner. See Figure S8.

Given the generalized effects of TTC1 depletion on mitochondrial health and integrity, we explored the possibility that these phenotypes may result from dysregulation of the mitochondrial proteome. We used tandem mass tag (TMT)-based quantitative proteomics to measure endogenous protein steady-state levels from whole-cell lysates of wild-type and TTC1-depleted K562 cells (Figure 5D), with the data shown here normalized to the mitochondrial proteome defined by the MitoCarta 3.0 database (Rath and Sharma et. al, 2021). We first noted a selectivity for the type of outer membrane protein showing a biogenesis defect, with β-barrel proteins such as the VDACs being slightly upregulated relative to other mitochondrial proteins (and remaining relatively unchanged overall; see also table S6) upon TTC1 depletion. In agreement with our reporter data, we also observed that endogenous CISD1 and MTCH2 levels were significantly impacted. Interestingly, we saw that SA protein TOMM20 which is a central receptor for import of all MTS proteins was the most significantly depleted and speculate that this could indirectly contribute to some of the TTC1-related mitochondrial health phenotypes. We further showed that TTC1’s impact on mitochondrial outer membrane substrates is likely post-transcriptional, as mRNA levels (assessed by qRT-PCR) of affected substrates were not significantly altered (Figure S8H). Finally, we established that the endogenous substrate levels were reproducibly altered in TTC1-depleted cells (Figure 5E) using immunoblotting.

TTC1 has a conserved TPR domain implicated in binding cytosolic HSP90 and HSP70 with a possible co-chaperoning function (Schuefler et. al, 2000, Lotz et. al, 2008). Both HSP90 and HSP70 have been implicated in aiding import of mitochondrial proteins (Deshaies et. al, 1988, Sheffield et. al, 1990, Young et. al, 2003, Fan et. al, 2006, Jores et. al, 2018). Therefore, while TTC1 could utilize HSP binding to aid SA protein targeting, we wanted to ensure that outer membrane biogenesis defects were not due to a loss of cytosolic HSP90 and/or HSP70. Indeed, we showed that HSP90 and HSP70 levels in whole-cell lysates were not impacted by TTC1 depletion, confirming that TTC1’s impact on outer membrane substrates is not an indirect consequence of cytosolic HSP loss (Figure 5E). Finally, to determine the extent of HSP binding by endogenous TTC1, we endogenously tagged all alleles of TTC1 with GFP in HEK293T cells and purified TTC1 with immunoprecipitation using an anti-GFP nanobody as previously described (Pleiner et. al, 2020). We determined interaction partners by separating proteins from the purified sample with SDS-PAGE followed by Coomassie staining and mass-spectrometry to identify discrete bands (Figure 5F). As a control for non-specific binding, we exogenously expressed and purified GFP from HEK293T cells. We saw that HSP90 (HSP90AA1, HSP90B) and HSP70 (HSPA1A, HSPA1B) were among the strongest TTC1-specific interactors. Additionally, we detected binding to a HSP70 co-chaperone protein BAG2 as well as to contact site proteins VSP13C and VAPB, consistent with prior observations (Lotz et. al, 2008). Due to TTC1’s general role in mitochondrial homeostasis and its ability to interact with cytosolic HSPs analogous to other TPR domain-containing membrane protein chaperones (i.e, TOMM70, SGTA; Young et. al, 2003, Philip et. al, 2013), we next sought to determine if TTC1 has an independent role in directly chaperoning and targeting nascent SA proteins to the outer membrane.

### TTC1 solubilizes and promotes nascent SA insertion in vitro, using a conserved C-terminal hydrophobic groove to direct substrate biogenesis

We posited that TTC1 depletion causes mitochondrial dysregulation by specifically impacting outer membrane protein biogenesis. Therefore, we sought to examine whether TTC1 can bind and directly target substrates to the OMM. For this, we utilized an in vitro system to directly probe the interactions between TTC1 and nascent SA substrates. To see whether TTC1 could specifically associate with mitochondrial SA proteins, we expressed flag-tagged TMD containing single-pass α-helical proteins in a rabbit reticulocyte lysate (RRL) system and assessed TTC1 interaction by native immunoprecipitation and immunoblotting. We saw an enrichment of TTC1 association with the SA proteins translated in RRL (CISD1, MIEF1, MTARC1, TOMM70) compared to ER TA protein SEC61β and inert globular VHP protein (Figure 6A). To understand the basis for these interactions, we interrogated the predicted structure and sequence conservation of TTC1 to determine functionally significant regions. Bioinformatic analyses across all sequence homologs of TTC1 (Landau et. al, 2005, Ashkenazy et. al, 2016, Yariv et. al, 2023) showed that the TPR and C-terminal domains of TTC1 are highly conserved (Figure 6B), while the N-terminal loop is poorly conserved and relatively unstructured (Figure S9A). Coloring of the AlphaFold2 (Jumper et. al, 2021) predicted model of TTC1 by electrostatic potential further indicates that the conserved C-terminal domain contains a highly hydrophobic groove, suggesting a possible interaction site with hydrophobic TMDs, analogous to other chaperones (Wang et. al, 2014, Mateja et. al, 2009, Meador et. al, 1992, Keenan et. al, 1998) (Figure 6B). Within the TPR domain, the ‘two-carboxylate’ clamp structure mediates electrostatic interactions with the C-terminal EEVD motifs of the HSP90 and/or HSP70 proteins (Schuefler et. al, 2000). These motifs are important in other proteins implicated in mitochondrial import, such as TOMM70. Indeed, directed alanine mutations in basic residues of the analogous TPR carboxylate clamp of TOMM70 reduce HSP binding and subsequent substrate mitochondrial import (Young et. al, 2003, Backes et. al, 2021). We thereby reasoned that this same motif may be critical to TTC1 function and tested the effects of putative HSP-binding mutants on substrate integration using the GFP11 reporter cassette in K562 IMS GFP1-10 cells with a flow cytometry readout of the resulting GFP:RFP ratios. We found that alanine mutations in the analogous TTC1 TPR residues (R190 and R194) partially impair the ability of exogenous TTC1 to rescue SA protein biogenesis relative to wild-type TTC1 (Figure 6C) without impacting TTC1 stability (Figure S9B). This was supported by a notable decrease in HSP90 binding for the TPR mutant compared to wild-type TTC1 (Figure S9C, left panel) which we assayed through co-immunoprecipitation of exogenous GFP-tagged TTC1 constructs as previously described.

**Figure 6.**
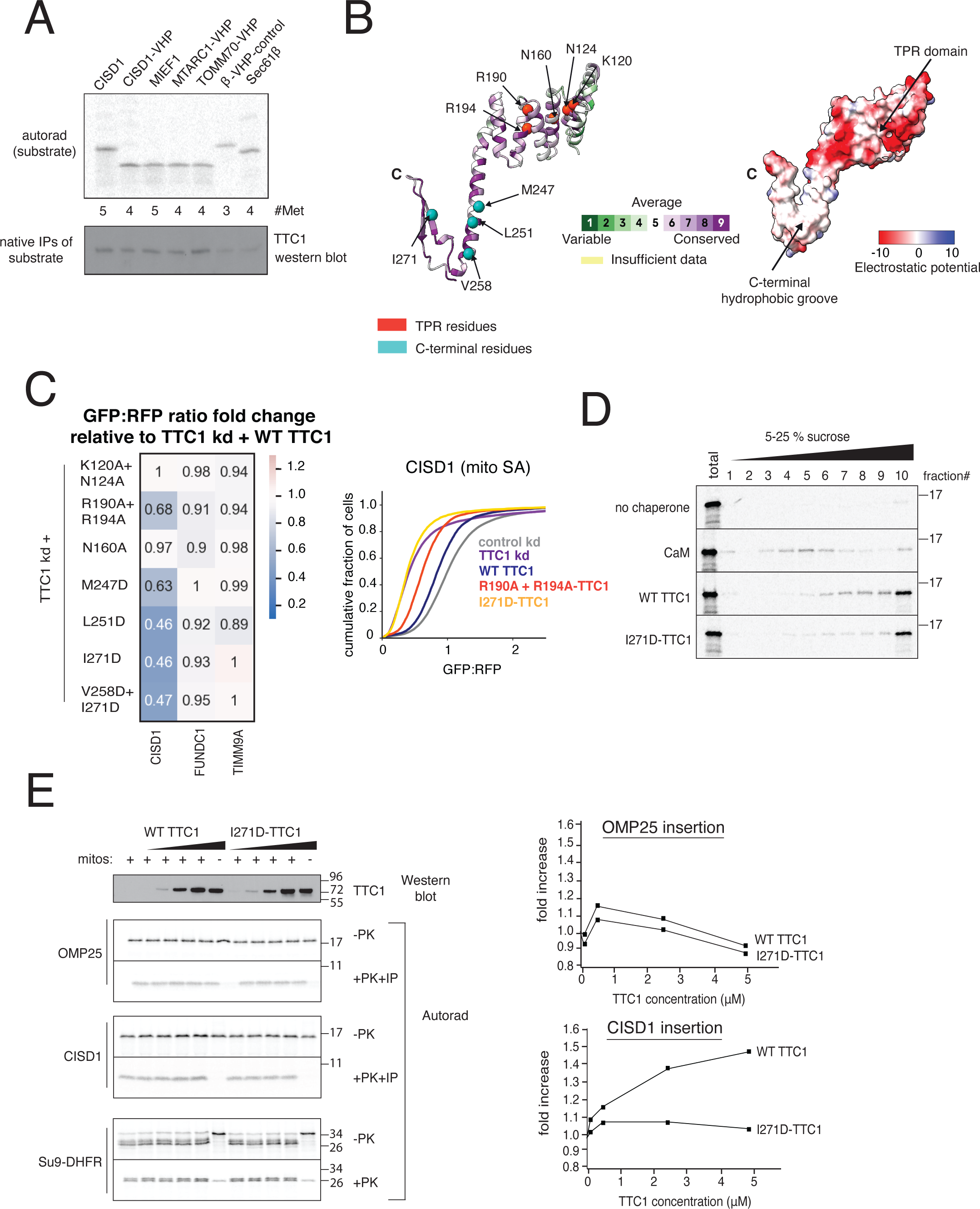
TTC1 chaperones nascent SA proteins and promotes insertion into mitochondria in vitro, using a C-terminal hydrophobic groove to direct biogenesis. (A) Indicated substrates tagged with 3xFLAG were translated in rabbit reticulocyte lysate (RRL) in the presence of ^35^S and released with puromycin. Immunoprecipitation and immunoblotting show the levels of TTC1 associated with the nascent substrates. (B) AlphaFold2 model of TTC1 structure colored by i) evolutionary conservation using the ConSurf server (see Methods for details) showing high conservation in the TPR domain and the C-terminus (left) and ii) electrostatic potential indicating a hydrophobic groove at the C-terminus (right). (C) The effects of putative heat-shock binding TPR domain mutants and C-terminal point mutants (individual residues indicated in (B)) on integration of indicated GFP11-fused reporters in K562 ZIM3 CRISPRi IMS GFP1-10 cells expressing a sgRNA targeting TTC1 were assessed in a similar manner to Figure 4D. (Left) Data is represented as a heatmap colored by the fold change in reporter integration (GFP:RFP ratio) for TTC1 depleted cells expressing WT or mutant TTC1. (Right) Flow data for the effects of the R190A+R194A TPR domain and I271D C-terminal mutants on SA reporter CISD1 is plotted as a cdf plot. (D) CISD1 was translated in vitro in the presence of ^35^S in the PURE system in the presence of either i) no chaperone, ii) purified calmodulin, iii) purified WT TTC1, iv) purified I271D-TTC1. The translations were run on a sucrose gradient and the extent of the substrate remaining soluble was determined by autoradiography. (D) (Left) Insertion into mitochondria isolated from wild-type K562 cells of endogenous CISD1 (SA), OMP25 (TA) and Su9-DHFR translated from RRL in the presence of increasing amounts of WT TTC1 or the I271D mutant (matched concentrations). Insertion is assessed by PK (proteinase K) digestion followed by an IP of the protected 6xHis tag. (Right) Quantification of the fold increase for mitochondrial insertion of OMP25 and CISD1 stimulated by WT and I271D-TTC1. See Figures S9 and S10.

To test the functional importance of the C-terminal domain of TTC1 and disrupt the hydrophobic surface, we introduced single or multiple aspartate mutations at the indicated residues (Figure 6B). We then assessed the abilities of these exogenous TTC1 mutants to rescue reporter biogenesis defects in K562 cells in the same manner as described for the TPR mutants. Strikingly, we observed that even single aspartate mutations in this groove completely abolished the biogenesis rescue by exogenous TTC1 (Figure 6C) without impacting protein stability (Figure S9B). These mutants did not exhibit diminished HSP90 binding (Figure S9C, right panel), arguing for a separation of function of the TPR and C-terminal domains in regulating substrate biogenesis. We further showed that the single I271D TTC1 mutant can moderately rescue some of the mitochondrial phenotypes caused by TTC1 depletion in cells, particularly network integrity (Figure S10A), and to a much lesser extent, certain parameters of mitochondrial respiration such as maximal respiration and ATP production (Figures S10B, C and D). This suggested that the C-terminal domain is more specifically involved in enabling substrate biogenesis rather than determining mitochondrial morphology.

To determine if TTC1 has any intrinsic TMD-chaperoning activity, we synthesized nascent CISD1 (representative SA substrate) in a chaperone-free *Escherichia coli*-based translation system assembled from purified translation factors (PURE system; Shimizu and Ueda, 2009) in the presence or absence of recombinant TTC1 (purified as shown in Figure S10E). As a control, we included Calmodulin (CaM), a factor known to recognize and shield hydrophobic domains in the cytosol (Shao and Hegde, 2011, Guna et. al, 2018). The extent of CISD1 remaining soluble was assessed by running the translations on a sucrose gradient. We first note that wild-type TTC1 was capable of preventing CISD1 aggregation, albeit to a slightly lesser extent than CaM (Figure 6D). Secondly, we show that this chaperoning activity is dependent on the C-terminal domain, as the I271D mutant resulted in significantly more aggregation of CISD1 (Figure 6D). Thus, we show that TTC1 can not only specifically associate with nascent signal-anchored proteins in the cytosol but also act as a chaperone independent of its HSP-binding co-chaperone function. Finally, we assessed the ability of purified wild-type and C-terminal mutant TTC1 proteins to stimulate insertion of substrates translated in RRL into wild-type mitochondria in vitro (Figure 6E). We saw that at matched concentrations, wild-type TTC1 but not the C-terminal mutant can specifically promote insertion of CISD1 into the mitochondria. In contrast, insertion of tail-anchored protein OMP25 or recombinant TOM substrate Su9-DHFR were unaffected by addition of TTC1 (Figure 6E). Cumulatively, the in vivo requirement of TTC1 for mitochondrial SA protein biogenesis and mitochondrial integrity, the physical engagement of substrates, and the ability to independently chaperone and promote insertion of the SA TMDS in vitro establish TTC1 as a novel cytosolic chaperone complex specifically required for α-helical OMM SA protein biogenesis.

## DISCUSSION

Here, we used a combination of systematic CRISPR-based genetic analyses, cell biological studies and biochemical characterization to delineate how mammalian cells ensure proper biogenesis of biophysically diverse α-helical OMM proteins. Our approach of studying multiple diverse substrates in a high-throughput manner allowed us to characterize cytosolic targeting factors previously not associated with OMM proteins and to uncover the rules governing α-helical outer membrane protein biogenesis in a principled manner. Taken together, our studies support a model (Fig. 7) for outer membrane α-helical protein biogenesis governed by three principles: i) Multiple parallel pathways are required to engage the full diversity of the α-helical OMM proteome, ii) Differential dependencies on biogenesis and quality control factors are defined by substrate type and topology, iii) Nascent substrates are triaged in the cytosol based on their type and topology using distinct chaperone networks. Specifically, polytopic and most SAs proteins rely on NAC and TTC1 cytosolic chaperone complexes respectively for outer membrane delivery whereas TA proteins depend on neither. NAC is strongly associated with the ribosome and known to engage nascent chains (Wiedmann et. al, 1994, Jomaa et. al, 2022, Lauring et. al, 1995, Gamerdinger et. al, 2015, Gamerdinger et. al, 2023). By contrast, TTC1 does not appear to associate with ribosomes and can engage SA substrates after release from the ribosome in vitro, suggesting that the primary mode of interaction is post-translational. While the cytosolic targeting factors triage the nascent α-helical proteins by type and topology, a wide range but not all biophysically diverse substrates converge at the MTCH1/2 insertases for integration at the membrane (Guna et. al, 2022). With regards to quality control, we observe a similar pattern, with the cytosolic quality control factor UBQLN1 differentiating substrates by type (only handling certain TA proteins) whereas the membrane-resident E3 ligase MARCHF5 impacts substrates of all types. More speculatively, TTC1’s association with BAG2 (Figure 5) could facilitate handoff to cytosolic E3 ligases for degradation as shown for ER TA proteins with BAG6 (Shao et. al, 2017). Finally, the alternate TOM complex receptor TOMM70 appears to coordinate biogenesis of polytopic and some TA proteins in cooperation with the MTCH1/2 insertases as has been implicated in yeast with the MIM insertases (Doan et. al, 2020). Overall, we present an outer membrane α-helical protein biogenesis model comprising multiple factors defined by triaging in the cytosol (both for targeting and quality control) based on substrate type and topology.

**Figure 7.**
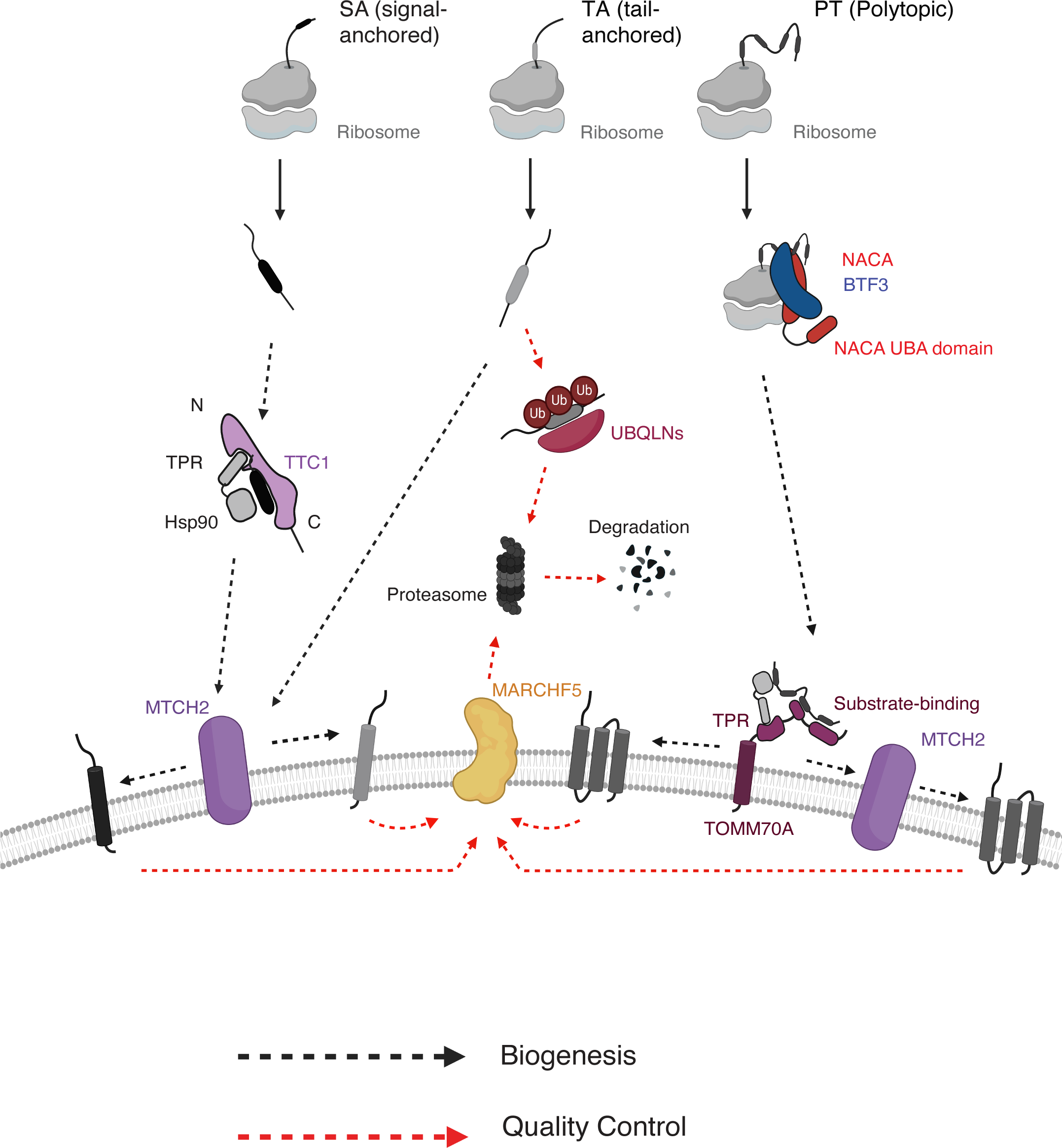
Working model for α-helical outer membrane protein biogenesis using multiple parallel pathways. Biogenesis pathways including cytosolic targeting, membrane insertion and quality control are depicted for three main types of α-helical outer membrane proteins (signal-anchored, tail-anchored and polytopic). Nascent outer membrane proteins are likely triaged during targeting in the cytosol, with signal-anchored proteins being targeted through cytosolic chaperone TTC1 and polytopic proteins targeted through co-translational chaperone complex NAC. Whether NAC directly engages and targets the polytopic proteins to the OMM remains to be explored. All proteins converge on MTCH1/2 insertases at the membrane for insertion, with some polytopic proteins depending more heavily on recruitment by TOMM70. Similarly, membrane-resident E3 ligase MARCHF5 handles quality control of aberrant/misfolded TMDs for substrates of all α-helical protein types, with some variation within protein type. In the cytosol, quality control processes are more divergent with tail-anchored proteins degraded using UBQLNs and no definitive pathway for other protein types.

Our observations that NAC depletion results in biogenesis defects for polytopic outer membrane proteins extend the role of NAC beyond canonical MTSs (Gamerdinger et. al, 2015, Jomaa et. al, 2022), as OMM proteins do not contain these MTSs or use the canonical TOM import machinery. NAC appears to promote direct outer membrane delivery of polytopic clients, rather than solely indirectly ensuring fidelity of mitochondrial targeting by preventing promiscuous SRP engagement and ER delivery which is the case for MTS-containing proteins (Wiedmann et. al, 1994, Lauring et. al, 1995, Gamerdinger et. al, 2015). Indeed, loss of NAC does not lead to increased delivery of OMM polytopic membrane proteins to the ER but rather their accumulation in the cytosol. This potential function of NAC in targeting of OMM polytopic α-helical proteins is analogous and distinct from the recently discovered role of NAC in facilitating direct targeting of ER signal sequences (Jomaa et. al, 2022). Positioned at the ribosome exit tunnel, NAC is poised to engage highly hydrophobic polytopic OMM proteins before they have a chance to aggregate. Whether the ultimate delivery to the OMM happens through NAC or a second mitochondrial-specific factor analogous to SRP for ER delivery is recruited to aid this process, and more generally how NAC orchestrates these decisions at the ribosome, remains to be explored.

By contrast, single-TMD α-helical OMM proteins do not require NAC for outer membrane delivery. However, they still need to be chaperoned in the cytosol and specifically targeted to the mitochondria, and post-translational targeting strategies for these and several other nuclear-encoded mitochondrial proteins have been discovered predominantly in yeast. Specifically, HSP chaperones and co-chaperones are known to be required for import of both outer membrane β-barrel (Jores et. al, 2018) and α-helical SA proteins (Drwesh et. al, 2022) in yeast. HSP70, HSP90 and HSP40 chaperone families have also been implicated in the import of MTS-containing proteins and inner membrane carrier proteins in yeast and human cells (Deshaies et. al, 1988, Young et. al, 2003, Hoseini et. al, 2016). However, the specific players in post-translational cytosolic chaperoning of outer membrane proteins in mammalian systems remained elusive. Here, we discover a cytosolic chaperone complex organized around TTC1, which is required for targeting α-helical SA proteins to the outer membrane. While TTC1 had not previously been implicated in mitochondrial function, we show that its depletion leads to specific defects in mitochondrial function with minimal impact on ER integrity. Like the MTCH1/2 insertases, TTC1 has no yeast homolog. TTC1 contains a conserved TPR domain with structural and sequence homology to other known TPR chaperones such as TOMM70, HOP and SGTA (Schuefler et. al, 2000). We show that TTC1 indeed uses the analogous HSP-binding dicarboxylate clamp implicated in TOMM70-directed substrate chaperoning and import (Young et. al, 2003, Backes et. al, 2021) to facilitate SA protein biogenesis. Importantly, TTC1 is distinguished from other TPR chaperones by the presence of a conserved C-terminal domain, which the high confidence alpha-fold model (Jumper et. al, 2021) predicts to form a hydrophobic groove. The integrity of this hydrophobic groove is essential for substrate biogenesis in vivo, as well as for TTC1’s ability to solubilize nascent SA proteins in vitro and promote their insertion into isolated mitochondria. Future work will be required to establish the exact mechanism by which this C-terminal hydrophobic groove contributes to TMD chaperoning and delivery. One possibility is that it provides a potential interaction site with hydrophobic TMDs, analogous to several other known chaperones (Wang et. al, 2014, Mateja et. al, 2009, Meador et. al, 1992, Keenan et. al, 1998). The detailed basis of substrate selection by TTC1 with regards to topology (i.e., its preferential engagement of SAs over TAs) also remains to be explored. It is possible that substrate hand-off to TTC1 depends on unknown machinery proximal to the ribosomal tunnel during synthesis.

The biogenesis model that emerges from our studies shows notable similarities in the logic used by the ER and the outer mitochondrial membrane to enable the cytosolic triaging of membrane proteins. Both biogenesis systems employ multiple distinct targeting pathways to the respective membranes, with substrates exploiting specific pathways by their type and topology. There appears to be a consistent requirement for polytopic membrane proteins to be co-translationally delivered to the membrane (in the ER through NAC and SRP, and in the mitochondria through NAC). Furthermore, single-pass substrates are chaperoned using similar strategies. For both mitochondrial and ER clients, the cell deploys distinct chaperone complexes such as TTC1 capable of recruiting HSPs to prevent substrate aggregation (SGTA for the ER; Cho and Shan, 2018) and containing conserved hydrophobic surfaces to facilitate TMD capture and delivery (Calmodulin and GET3 for the ER). One apparent difference between the two membrane biogenesis systems concerns the variety of insertion machinery at the membrane – OMM α-helical proteins widely depend on the MTCH1/2 insertases, while ER membrane proteins utilize the SEC61 translocon and the EMC and GET complexes, as well as TMD assembly factors such as CCDC47 (Chitwood et. al, 2020, Ast et. al, 2013, Aviram et. al, 2016, Costa et. al, 2018). However, a subset of OMM α-helical proteins show comparatively weaker dependencies on the MTCH1/2 insertases (such as SA protein USP30 and polytopic protein TSPO) for biogenesis, leading us to speculate that there may be yet undefined machinery recruiting and inserting these proteins at the OMM which could have been missed due to genetic redundancy.

Overall, our systematic discoveries of the parallel pathways and auxiliary proteins involved emphasize the complexity and nuances of outer mitochondrial membrane α-helical protein biogenesis in mammals. These systems appear to have evolved to handle biogenesis of a larger and more complex α-helical OMM proteome with TTC1 and MTCH1/2, both of which are absent in the analogous yeast systems. The existence of multiple distinct biogenesis and quality control factors with a high degree of substrate specificity could allow individual mitochondria to regulate their outer membrane composition and thereby communication with the cytosol and other organelles under various physiological conditions and disease states. We anticipate that these discoveries could be adapted to tune the outer mitochondrial membrane proteome at high precision, which in turn could inform efforts to mitigate outer membrane related diseases such as Alzheimer’s, Parkinson’s and a variety of cancers (Bose and Beal, 2016, Vyas et. al, 2016, Wang et. al, 2020(2)).

## ACKNOWLEDGEMENTS

We thank J. Nunnari and M. Le Vasseur for sharing the mitochondrial split GFP system, and J.M. Replogle for helping establish the CRISPRi screening system using the split GFP strategy. We thank Y.H. Chen for assistance with computational analyses, and Z. Levine and K.E. Yost for careful reading and input on the manuscript. We thank the Whitehead Institute and Koch Institute Flow Cytometry Cores for access to FACS machines; the Whitehead Institute Genome Technology Core for support with sequencing of screen libraries; the Whitehead Institute W.M. Keck Microscopy Facility for access to confocal microscopes and support with imaging analysis; and the Whitehead Proteomics Core for support for mass spectrometry experiments. This work was supported by the NIH pre-doctoral training grant T32GM007287 (G.M.), the Larry L. Hillblom Foundation (A.J.I.), the Fannie and John Hertz Foundation and NSF Graduate Research Fellowships (R.A.S.), the Heritage Medical Research Institute (R.M.V.) Human Frontier Science Program 2019L/LT000858 (A.G.), and the Howard Hughes Medical Institute (J.S.W.).

## Author contributions

G.M., A.G. and J.S.W. were responsible for the conception, design and interpretation of experiments and wrote the manuscript with input from all authors. G.M. led all in cell experiments and accompanying data analysis with help from T.K.E., who performed all qRT-PCR experiments and assisted with flow cytometry and immunoprecipitation studies. T.A.S., A.J.I., and A.G led all in vitro experiments and performed accompanying data analysis, with supervision from R.M.V. R.A.S. performed analysis of genome-scale perturb-seq data and cloned plasmids pRS447 and pRS508. F.S. led all mass spectrometry experiments and accompanying data analysis. J.S.W. and R.M.V. funded the study.

## Data and materials availability

All data needed to evaluate the conclusions in this paper are present in the paper or the supplementary materials.

## MATERIALS AND METHODS

### Materials/Reagents

**Antibodies and Reagents**: see Table S9.

### Cell culture and lentiviral production

K562 cells were grown in RPMI-1640 containing 25mM HEPES, 2.0 g/L NaHCO3, 0.3 g/L L-Glutamine supplemented with 10% FBS, 2 mM glutamine, 100 units/mL penicillin and 100 mg/mL streptomycin. Cells were maintained between 0.25 x 10^6^ -1 x 10^6^/ml. HEK293T cells were grown in Dulbecco’s modified eagle medium (DMEM) containing 10% FBS, 2 mM glutamine, 100 units/mL penicillin and 100 mg/mL streptomycin. hTERT-immortalized RPE1 cells were grown in DMEM:F12 supplemented with 10% FBS, 2 mM glutamine, 100 units/mL penicillin and 100 mg/mL streptomycin. All cell lines were grown at 37°C and 5% CO_2_. To produce lentivirus, HEK293T cells were co-transfected with transfer plasmids (pCMV-VSV-G and delta8.9, Addgene #8454) and standard packaging vectors using the TransIT-LT1 Transfection Reagent (Mirus, MIR 2306). Supernatant was collected 48 hours post transfection, centrifuged, aliquoted and flash frozen at -80C until further use. Virus for the genome-wide CRISPRi screens was also generated using this method with one exception – HEK293T cells were seeded in IMDM (Thermo Fisher Scientific #1244053) supplemented with 20% inactivated fetal bovine serum (GeminiBio #100-106). In all instances, virus was rapidly thawed prior to transfection.

### Cell line generation

CRISPRi cell lines expressing GFP1-10 targeted to the IMS were made using either the MICU1 or LACTB targeting seqeunces as previously described – resulting in the K562 CRISPRi lines with MICU1 GFP1-10 or LACTB GFP 1-10 in either KOX1 or ZIM3 KRAB domain containing cells (Guna et. al, 2022). To monitor reporter integration into the mitochondrial matrix, a K562 cell line was similarly generated by infecting K562 CRISPRi cells expressing dCas9-BFP-KOX1-KRAB with lentivirus containing the COX4 GFP1-10 construct (Gilbert et. al, 2014). Post infection, cells were sorted as single-cell clones in 96-well plates using a Sony Cell Sorter (SH800S) and confirmed after expansion by complementation with a COX4-GFP11 construct.

For use in microscopy studies, we generated RPE1 cell lines. In all cases, RPE1 cells expressing ZIM3 KRAB-dCas9-P2A-BFP were used, as previously described (Replogle and Saunders et. al, 2022). To monitor split-GFP reporter integration into the OMM (Figure S1A), RPE1 ZIM3 CRISPRi cells were transfected with virus containing the MICU1 GFP1-10 construct, single-cell sorted using the Sony Cell Sorter and confirmed by complementation with the MICU1-GFP11 construct.

Cell lines for the genome-wide screens were generated by co-infecting virus containing MICU1(GFP1-10) and RFP-P2A-(reporter)-GFP11 (GFP11-CISD1, FUNDC1-GFP11, GFP11-TSPO) under a tet-inducible promoter to one copy per cell in K562-dCas9-BFP-KRAB Tet-on cells (Jost et. al, 2017). Post infection, cells were sorted into single-cell clones, expanded and confirmed by induction with doxycycline at 100ng/ml as well as microscopy with MitoTracker DeepRed dye to confirm predominantly mitochondrial localization of the reporters.

To generate the TTC1-GFP cell line used for immunoprecipitation (Figure 5F), HEK293T cells were transfected with SpCas9+sgRNA (pX330 backbone) and HDR donor plasmids at a 1:1 ratio with Opti-MEM reduced serum media and TransIT-LT1 Transfection Reagent. 6 days post-transfection, cells were sorted for GFP (single clones) using the Sony Cell Sorter and expanded. Clones were validated using PCR amplification with primers outside the endogenous 5’ and 3’ homology arms and sequencing, as well as western blots to confirm tagging at all loci.

### Constructs

Endogenous sequences used in this study for in vitro and in vivo experiments were sourced from UniProtKB/Swiss-Prot and included: CDGSH iron-sulfur domain-containing protein 1 (CISD1, **Q9NZ45**); synaptojanin-2 binding protein (OMP25/SYNJBP; **P57105-1**), FUN14 domain-containing protein 1 (FUNDC1; **Q8IVP5-1**), translocator protein (TSPO; **P30536)**, mitochondrial amidoxime-reducing component 1 (MTARC1; **Q5VT66)**, mitochondrial dynamics protein MID51 (MIEF1; **Q9NQG6**), mitochondrial import receptor subunit TOM20 homolog (TOM20; **Q15388**), ubiquitin carboxyl-terminal hydrolase 30 (USP30; **Q70CQ3-1**), mitochondrial import receptor subunit TOM70 (TOMM70; **O94826**), mitochondrial cardiolipin hydrolase(PLD6; **Q8N2A8**), mitochondrial antiviral-signaling protein (MAVS; **Q7Z434-1**), mitochondrial Rho GTPase 1 (RHOT1; **Q8IXI2-1**), Bcl2-like protein 1 (BCL2L1; **Q07817-1**), mitochondrial import receptor subunit TOM22 homolog (TOM22; **Q9NS69-1**), E3 ubiquitin-protein ligase MARCHF5 (MARCHF5; **Q9NX47**), mitochondrial ubiquitin ligase activator of NFKB 1 (MUL1; **Q969V5**), mitochondrial carrier homolog 2 (MTCH2; **Q9Y6C9**), mitochondrial import inner membrane translocase subunit Tim9 (TIMM9; **Q9Y5J7-1**), mitochondrial import inner membrane translocase subunit Tim10 (TIMM10; **P62072**), cytochrome c oxidase subunit 4 isoform 1, mitochondrial (COX4I1; **P13073**), 60 kDa heat shock protein, mitochondrial (HSPD1; **P10809**), squalene synthase isoform 1 (SQS/FDFT1; **Q6IAX1**), vesicle associated membrane protein 2 (VAMP; **P51809-1**), protein transport protein Sec61 subunit beta (SEC61B; **P60468**), alkaline ceramidase 1 (ACER1; **Q8TDN7**), calreticulin (CALR; **P27797**), nascent polypeptide-associated complex subunit alpha (NACA; **Q13765**), transcription factor BTF3 (BTF3; **P20290**), and tetratricopeptide repeat protein 1 (TTC1; **Q99614**). All sequences were cloned into in vivo expression vectors using gene blocks synthesized by IDT (Integrated DNA Technologies, Coralville, IA).

Constructs for expression in the RRL system were based on the SP64 vector (Promega) while those for expression in the PURE system (Shimizu and Ueda, 2009) were based on the T7 PURExpress plasmid provided by New England Biolabs. For in vitro reactions, an N-terminal 3XFlag tag and a C-terminal 6x tag were appended to either full length endogenous proteins or the TMD with flanking regions to allow for purification as previously described (Guna et. al, 2022). Where indicated, the villin headpiece (VHP) domain was conjugated to the indicated termini. The TOM dependent substrate Su9-DHFR was designed by fusing dihydrofolate reductase (DHFR) to the mitochondrial transit sequence from *N. crassa* ATP synthase subunit 9 (residues 1–69).

All constructs for expression in K562 cells were cloned in a mammalian expression lenti-viral backbone containing a UCOE-EF-1α promoter and a 3′ WPRE element ((J.J. Chen et. al, 2019); Addgene #135448) by Gibson assembly. Dual fluorescent reporters used for the CRISPRi screens (RFP-P2A-GFP11-CISD1, RFP-P2A-FUNDC1-GFP11, RFP-P2A-GFP11-TSPO and RFP-P2A-OMP25-GFP11) were integrated into an SFFV-tet3G backbone cloning (M. Jost et. al, 2017) to allow for inducible expression. The dual color reporter system used for in vivo flow cytometry experiments has been previously described (Chitwood et. al, 2018, Guna et. al, 2018). For simpler nomenclature, the mCherry variant of RFP, used throughout the manuscript, is annotated as ‘RFP’. The GFP11 tag (RDHMVLHEYVNAAGIT) was appended to the appropriate termini of the proteins as determined by predicted topology to allow for GFP 1-10 complementation (see Figure 1A). For targeting GFP 1-10 to the ER lumen, the GFP 1-10 KDEL was preceded by the human calreticulin signal sequence (Cabantous et. al, 2005, Kamiyama et. al, 2016, Inglis et. al, 2020). For targeting to the mitochondrial intermembrane space, GFP 1-10 was preceded by either the targeting sequence from MICU1 (residues 1–60) (Le Vasseur et. al, 2021, Kamiyama et. al, 2016) or that of LACTB (residues 1–68) (Hung et. al, 2014). Finally, for targeting to the mitochondrial matrix, GFP1-10 was preceded by the targeting sequence from CoxIV – Cox protein targeting sequences have been previously described for targeting to the matrix (Le Vasseur et. al, 2021). For in cell rescue experiments, expression of MTCH2 was from a BFP-P2A-(MTCH2) cassette (Guna et. al, 2022). Exogenous TTC1 and TOMM70 proteins were expressed using a similar vector, with the addition of an N-terminal 3xFLAG tag (BFP-P2A-3xFLAG-(TTC1 or TOMM70) cassette). Point mutations for TTC1 constructs were made with primers designed using the New England Biolabs Q5 Site-Directed Mutagenesis Kit – these were K120A+N124A, R190A+R194A, N160A, M247D, L251D, I271D, and V258D+I1271D. NACA and BTF3 dual expression vectors were designed within a BFP-P2A-3xFLAG-(NACA)-T2A-6xHis-(BTF3) cassette; point mutations were K78E and D205R+N208R for NACA, and K43E for NACB. For cell GFP-based immunoprecipitation experiments, TTC1 and associated mutants were cloned in a BFP-P2A-GFP-(TTC1) cassette by Gibson assembly.

Single sgRNA constructs were generated by annealed oligo cloning of top and bottom oligonucleotides synthesized by IDT (for all protospacer sequences used, see table S8) into a lentiviral pU6-sgRNA EF-1α-Puro-T2A-BFP vector digested with BstXI/BlpI (Addgene, #84832). The exception was the sgRNAs for TTC1 and Control used in the Seahorse assay, with the annealed oligos cloned into a lentiviral pU6-sgRNA EF1 -1α-Puro-T2A-GFP vector digested with BstXI/BlpI (Replogle and Saunders et. al, 2022). In guides used for rescue experiments, the BFP was excised to prevent fluorophore interference. To deplete multiple genes at once (see Figures 3 and 4) or to increase knock-down efficiency, we used a programmed dual sgRNA guide vector (Addgene #140096) (see table S8 for list of dual guide combinations and respective protospacer sequences). Tagging endogenous TTC1 in HEK293T cells with GFP was performed as previously described (Zhang et. al, 2017). The sgRNA was generated by annealed oligo cloning into a chimeric human codon-optimized SpCas9 and pU6-driven guide RNA expression plasmid (pX330, Addgene #42230) with BbsI digestion (see table S8 for protospacer sequence). The donor plasmid was cloned into the empty backbone pUC19 cloning vector (Addgene #50005) digested with SaII and KpnI – the 5’ and 3’ homology arms at the TTC1 C-terminus appended with the TTC1 protospacer + PAM sequence on both ends, and the 20xGS-linker + sfGFP insert were amplified from IDT-synthesized gene blocks and cloned into the digested backbone by Gibson assembly.

All constructs for bacterial protein expression were cloned in the pQE plasmid (Qiagen, Valencia, CA). WT and mutant TTC1 were cloned downstream of a His_14_-*bd*SUMO-GFP tag. The construct for the expression of bdSENP1(Addgene #104962; Frey and Görlich et. al, 2014) and CaM have been previously described (Shao et. al, 2017).

All plasmids are available upon request and described in table S8.

### CRISPRi screens

Genome-scale FACS-based CRISPRi screens were performed as previously described (Gilbert et. al, 2014, Horlbeck et. al, 2016, Guna et. al, 2022). All screens were performed in duplicate. For each, the hCRISPRi-V2 compact library (Addgene #83969, 5 sgRNAs/TSS) was transduced at a MOI (multiplicity of infection <1) into 350 million cells. The percentage of cells transduced was measured at 48 hours post transduction by recording the percent of BFP positive cells (∼20-30%). Replicates were grown separately in 1L of RPMI-1640 in 3L spinner flasks (Bellco, SKU: 1965-61030). The cells were selected with 1 ug/ml puromycin starting at 48 hours post-transduction for 3 days until the transduced cells accounted for 90-95% of the population, and then maintained at 0.5 x 10^6^/ml to ensure an average coverage of more than 1000 cells/sgRNA. 36 hours prior to sorting, expression of the reporters was induced with doxycycline at 100 ng/ml. For sorting on a BD FACS Aria II, cells were gated for BFP (indicating guide-positive cells) and on the GFP:RFP ratio (highest and lowest 30%). Approximately 40 million cells each were collected from the top and bottom 30% based on the GFP:RFP ratio, pelleted and flash frozen. Genomic DNA was isolated from frozen cells using the Nucleospin Blood XL kit (Takara Bio, #740950.10) and amplified with barcoded primers by index PCR (see table S10 for sequences). Guide libraries (∼264 bp) were purified using SPRIbeads (SPRIselect Beckman Coulter #B23318) followed by sequencing with an Illumina HiSeq2500 high throughput sequencer.

### Processing of CRISPRi screen data

Using the Python-based ScreenProcessing pipeline, sequencing reads were aligned to the CRISPRi v2 library sequences, counted and quantified (https://github.com/mhorlbeck/ScreenProcessing, Horlbeck et. al, 2016). Negative control gene generation and calculation of phenotypes and Mann-Whitney p values was performed as previously described (Gilbert et. al, 2014; Horlbeck et. al, 2016). Phenotypes from sgRNAs targeting the same gene were collapsed into a single GFP:RFP ratio phenotype derived from the average of the top three scoring sgRNAs (by absolute value) and assigned a p-value through the Mann-Whitney test of all sgRNAs targeting the same gene compared to the non-targeting controls. All additional CRISPRi screen data analyses were performed in Python 3.6.9 using a combination of Numpy (v1.19.0), Pandas (v0.25.0), and Scipy (v1.5.4). Volcano plots were generated using the Python Bokeh package (v 3.1.1). Gene-level phenotypes and counts are available in tables S1-S4.

### Flow cytometry

For all reporter experiments, constructs were individually packaged into lentivirus, spinfected into K562 cells and analyzed by flow cytometry after 48-72 hours. For experiments testing the effects of single or dual sgRNAs on reporters, sgRNA packaged into lentivirus was first spinfected into K562 cells at a MOI <1 (∼20-40%). Transduced cells were selected with puromycin at 1 ug/ml for 3 days starting at 48 hours post-spinfection, subsequently spinfected with reporter virus at 48 hours post puromycin-recovery and analyzed 48-72 hours after. With experiments testing rescue by adding back exogenous wild-type or mutant factors, virus was first titered by flow cytometry to match expression levels. Titered factor virus and reporter virus were co-transduced into guide-selected K562 cells at 24 hours post puromycin recovery and analyzed by flow cytometry 72 hours after. Where indicated, cells were sorted by % BFP-positive concurrently with flow cytometry analysis to select for cells either expressing i) guides, ii) titered rescue factors, for downstream qRT-PCR and immunoblotting analyses. All flow cytometry data was collected on an NXT Flow Cytometer (Thermo Fisher). Data analysis was performed in Python using the FlowCytometryTools package and a combination of Pandas, Numpy, Scipy and Seaborn libraries.

### qRT-PCR

Cells treated with sgRNAs of interest were washed with PBS, pelleted and flash frozen. Total RNA was isolated from frozen cells using the RNeasy Mini kit (74104, Qiagen). Cells were resuspended as per manufacturer’s instructions for RNAse-rich cell lines using 2-MercaptoEthanol (Sigma-Aldrich, M6250-250 mL) and on-column DNAse digestion was performed to further purify the product (Qiagen, 79254). Isolated RNA was converted to cDNA using the Maxima First Strand cDNA Synthesis Kit for qRT-PCR as per manufacturer’s instructions (Thermo Fisher, K1641). Reactions were performed with the Dynamo ColorFlash SYBR Green qPCR kit (Thermo Fisher, F416L) per manufacturer’s instructions on a QuantStudio 7 Flex Real-Time PCR system (Thermo Fisher, 4485701). The relative expression ratios were calculated relative to the housekeeping gene GAPDH using the 2^-ΔΔCt^ method. Experiments were performed in technical triplicate with means and standard deviations plotted. Primers used are included in table S10.

### SDS-PAGE and western blot

Cells (approx. 0.5 - 1x 10^6^ per sample) were first washed with PBS and pelleted, and then lysed in buffer containing 50mM HEPES pH 7.5, 200mM NaCl_2_, 2mM MgAc_2_, 1% Triton X-100 and 1x Pierce complete protease and phosphatase inhibitor cocktail (Thermo Fisher, A32959). Lysates were clarified to remove cell debris by rotating end-over-end at 4°C briefly and subsequently centrifugated at 5,000g for 5 minutes at 4°C. Total protein material for each sample was quantified using the Pierce BCA Protein Assay kit (Thermo Fisher, 23225) for equal loading. Samples were first boiled with 6x Laemmli SDS buffer at 95°C for 5 minutes and then proteins were separated on Bolt 4-12% Bis-Tris gels (Thermo Fisher, NWO4122BOX and NWO4127BOX), transferred to Nitrocellulose membranes using the Mini Trans-Blot Cell (BioRad) kit at Mixed MW (molecular weight) settings according to the manufacturer’s instructions, blocked with EveryBlot Blocking Buffer (BioRad 12010020) at room temperature for 1 hour, and subsequently probed. For a list of primary antibodies used, see table S9. Secondary antibodies used were goat anti-mouse- and anti-rabbit-HRP (#172-1011 and #170-6515, BioRad, USA).

### Live-cell confocal microscopy

Reporters were packaged individually as lentivirus and transduced into RPE1 cells at a MOI<1. Cells expressing reporters (Figures S1A and S6B) were selected by sorting for the relevant fluorophores (S1A: RFP, S6B: GFP) on the Sony Cell Sorter and subsequently plated in 35 mm-high µ-dishes with a polymer coverslip bottom at a density of approximately 0.5 - 1 x 10^5^ cells/dish (Ibidi, 81156). 15 minutes prior to imaging, cells were incubated with DMEM:F12 containing MitoTracker DeepRed FM at a concentration of 75 nM (Thermo Fisher, M22426) to visualize the mitochondria at 37°C. Cells were subsequently imaged on a Zeiss LSM 980 with Airyscan2 Laser Scanning Confocal microscope with incubation at 37°C and humidified CO_2_ with 63x magnification, 1.4 numerical aperture.

### Cell fixation, immunofluorescence and confocal microscopy

For immunofluorescence experiments (Figures 4B, 5B and S6C-F) and experiments studying mitochondrial morphology (Figures 5A, S8F-G and S10A), RPE1 cells were selected for expressing the indicated sgRNAs with puromycin at 5 ug/ml, recovered and plated in 35 mm-high µ-dishes with a polymer coverslip bottom at a density of approximately 0.5 - 1 x 10^5^ cells/dish (Ibidi, 81156) 7 days post guide-transduction. To visualize mitochondria, cells were incubated with DMEM F:12 containing MitoTracker DeepRed FM at a concentration of 150 nM (Thermo Fisher, M22426) at 37°C for 45 minutes. Cells were then rinsed once in complete medium, once in ice-cold 1x PBS and then fixed with 4% paraformaldehyde in PBS pH 7.4 for 10 minutes at room temperature. Post-fixation, cells were washed three times with ice-cold 1x PBS and permeabilized in 0.1% triton X-100 in PBS for 5 minutes at room temperature. Cells were then blocked with 10% normal goat serum (NGS) (Abcam, ab7481) in PBS at room temperature for 1 hour. Primary antibodies were added at the following dilutions in 1% BSA (bovine serum albumin) in PBS: rabbit anti-TSPO and mouse anti-PDI at 1:500, rabbit anti-Sec61β and rabbit anti-MAVS at 1:200 for 1 hour at room temperature. Cells were then washed three times with ice-cold 1x PBS for 5 minutes each and incubated with Alexa-Fluor-conjugated secondary antibodies diluted in 1x PBS at 1:1000 for 1 hour at room temperature. Cells were finally washed three times with ice-cold 1x PBS for 5 minutes each and mounted with mounting media containing DAPI (Ibidi, 50011). Imaging was performed on a Zeiss LSM 980 with Airyscan2 Laser Scanning Confocal microscope with 63x magnification, 1.4 numerical aperture.

### Seahorse assays

K562 cells expressing ZIM3 dCas9-BFP-KRAB were spinfected with GFP-tagged CRISPRi guide plasmids (Replogle and Saunders et. al, 2022) for the seahorse assays in Figures 5C and S10B-C. 48 hours post spinfection, cells were treated with puromycin for three days at 1 ug/ml and allowed to recover for 24 hours. Subsequently, cells were spinfected with lentivirus carrying i) BFP alone (control), ii) BFP and exogenous WT TTC1, iii) BFP and exogenous mutant I271D-TTC1. Cells were then sorted on the basis of GFP (guide-containing) and BFP (rescue constructs), at 48 hours post spinfection. The following day, seahorse assay plates were treated with Cell-Tak (Corning 354240) to help adhere K562 cells according to the manufacturer’s instructions. Cells were collected by centrifugation and resuspended in supplemented Seahorse XF RPMI (Agilent 103576-100) at ∼8 x 10^5^/ml. 40,000 cells were added to each well on the Seahorse assay plate with 10 wells/condition and attached via centrifugation at 200xg for 1 minute. Post recovery for 30 mins at 37°C, the cells were subjected to a Mito Stress Test on a Seahorse XFe96 analyzer according to the manufacturer’s instructions. From the same batch of sorted cells used for the Seahorse experiment, cells were pelleted, washed with PBS, lysed and assayed for expression of TTC1 rescue constructs by SDS-PAGE and immunoblotting as described above.

### Mass spectrometry

### Extraction and tryptic digestion of proteins from whole cells and IP-affinity enrichments

Proteins from whole cell extracts and IP-affinity enrichments were isolated using the S-trap protocol. (Shevchenko et. al, 2007, Rappsilber et. al, 2007, Schulte et. al, 2019). The fractions were lysed by adding 100 µL S-trap lysis buffer (5 % (w/v) SDS, 50 mM TEAB pH 8.5) followed by an incubation step for 10 min at 70°C. To prepare proteins isolated via IP-affinity purification, 10 µL raw extract in IP-buffer (50mM HEPES pH 7.5, 200mM NaCl_2_, 2mM MgAc_2_, 1 mM DTT, 1% triton x-100 and 1x Pierce complete protease and phosphatase inhibitor cocktail (Thermo Fisher, A32959)) were mixed with 10 µL 2x S-trap Lysis buffer (10 % (w/v) SDS, 100 mM TEAB pH 8.5) and heated for 10 min at 70 °C (Shevchenko et. al, 2007). Protein raw extracts were centrifuged for 10 min at 20,000 x g and the clarified solutions were transferred to fresh tubes. Protein concentrations were determined using the Pierce 660 nm Protein Assay Reagent together with the Ionic Detergent Compatibility Reagent (Thermo Scientific, Waltham, MA).

For the purification of proteins via S-trap (Profiti, Fairport, NY, USA), 50 µg protein or 20 µL IP-affinity extracts were diluted in a total volume of 23 µL S-trap lysis buffer. Cysteines were reduced with 1 µl 250 mM TCEP followed by an alkylation step with 5 µL 200 mM MMTS by letting each of the reactions incubate for 15 min at room temperature, respectively. For protein trapping, the extracts were acidified with 2.5 µL 27.5 % (w/v) phosphoric acid and dissolved proteins were precipitated by adding 165 µL S-trap binding buffer (100 mM TEAB pH 7.6 in 90 % (v/v) ice-cold methanol). The total volume was applied to the S-trap Micro columns and the precipitated proteins were separated from suspension via centrifugation for 1 min at 4000 x g. Salts and detergents were removed by washing the S-trap columns six times with 150 µL S-trap binding buffer. Next the columns were placed into fresh Low Protein Binding Microcentrifuge Tubes (Thermo Scientific, Waltham, MA, USA) to harvest tryptic peptides after digestion. Proteins caught by the S-trap fiber glass filters were enzymatically digested by adding 30 µL 1 µg trypsin/LysC mix dissolved 50 mM TEAB pH 8.5 and letting the reactions incubate overnight at 37°C in a humidified incubator. Tryptic peptides dissolved in trypsin/LysC buffer were collected by centrifuging the columns for 1 min at 4000 x g followed by three washes with 40 µL 50 mM TEAB pH 8.5, 40 µL 0.2 % (v/v) formic acid in water, and 40 µL 50 % acetonitrile in water, to wash off peptides remaining on the fiberglass filter and the C18 plug of the S-traps. The peptides were pooled, lyophilized, and dissolved in 20 µL 100 mM TEAB pH 8.5 and stored at -20°C until being used for TMT labeling. The IP digests were dissolved in 20 µL 0.2 % (v/v) formic acid in water and were used for LC-MS measurements without any further sample preparation steps.

### TMT-labeling, solid phase extraction, and peptide fractionation

Peptide concentrations of whole cell S-trap extracts in 100 mM TEAB were quantified using the Pierce Quantitative Fluorometric Peptide Assay (Thermo Scientific, Waltham, MA, USA) according to the manufacturer’s instructions. For each labeling reaction, 10 µg peptide was diluted in 75 µL 100 mM TEAB pH 8.5, mixed with 25 µL acetonitrile, and 200 µg TMT 6plex label (Thermo Scientific, Waltham, MA, USA) dissolved in 5 µL anhydrous acetonitrile was added. The labeling reactions were vortexed immediately, incubated for 60 min at room temperature, and remaining TMT reagents were subsequently inactivated with 2 µL 5 % (w/v) hydroxylamine for 30 min. After quenching, samples were pooled, lyophilized, and resuspended in 300 µL 0.2 % TFA in water. Resuspended isotope-labeled peptides were loaded onto Pierce High pH Reversed-Phase Peptide Fractionation columns (Waltham, MA, USA) followed by washing and elution using the manufacturer’s recommendations for the fractionation of TMT-labeled samples. The individual peptide fractions were lyophilized and reconstituted in 20 µL 0.2 % formic acid in water and were employed for LC-MS analysis.

### LC-Orbitrap-MS

LC-MS/MS data was acquired using an EasyLC 1200 nanoLC operated together with an Orbitrap Exploris mass spectrometer and a FAIMS Pro ESI source unit (Thermo Fisher Scientific, Waltham, MA, USA). The injection volume was 1 µL for TMT-labeled peptides and 5 µL for IP extracts, and peptides were separated using a 25 cm EasySpray nanoLC column (Thermo Fisher Scientific ES902) protected by a 2 cm Acclaim PepMap 100 guard column (Thermo Fisher Scientific 164946). The mobile phases 0.1 % (v/v) formic acid in water (solvent A) and 0.1 % (v/v) formic acid in 80 % (v/v) acetonitrile (solvent B) were utilized for chromatographic separation. The gradient parameters were as follows for TMT-labeled peptide fractions: 0.3 µL/min flow rate, 40°C column temperature, a gradient from 5 % solvent B up to 40 % solvent B over 120 min for peptide separation, and 10 min 95% solvent B as well as a see-saw gradient oscillating between 98 and 2 % solvent B in 3 x 2 min steps for the removal of remaining peptides. For IP digests, the following settings were employed: 0.3 µL/min flow rate, 40 °C column temperature, a gradient from 2 % solvent B up to 40 % solvent B over 48 min for peptide separation, and 12 min 100% solvent B as well as a see-saw gradient oscillating between 98 and 2 % solvent B in 1 x 3 min steps for the removal of remaining peptides.

The ion source temperature was maintained at 270 °C with a voltage of 1800 V (positive mode) and all ions generated by the ESI source were filtered by the FAIMS Pro unit at -50 and -65 V. The MS1 for TMT-labeled samples parameters were as follows: 120,000 resolution in DDA (data-dependent acquisition) mode, m/z 350-1200 mass range, automatic gain control (AGC) target, and automatic maximum injection time. For IP digests, 60,000 resolution in DDA mode across a m/z range of 350-1400 were utilized with all other parameters remaining unchanged.

The MS/MS was measured using the following specifications in TMT experiments: pick all precursor ions above 5 x 10^3^ intensity, charge state +2 to +5, isolation window of m/z 0.7, high energy collisional dissociation (HCD) at a normalized collision energy of 36, precursor ion exclusion for 60 sec. Resulting MS2 product ions were analyzed with automatic AGC target setting and automatic maximum ion accumulation time in the Orbitrap mass analyzer at 30,000 resolution. For IP peptides, ions with a charge state of +2 to +6 were selected within an isolation window of m/z 1.3 and a collision energy of 30 at 15,000 resolution while maintaining the other settings.

### Immunoprecipitation and detection of binding partners

In order to detect interaction partners of TTC1, endogenous GFP-tagged TTC1 was isolated from cells using a previously described GFP purification technique (Pleiner et. al, 2020), and interaction partners were probed using SDS-PAGE separation, Coomassie staining and mass spectrometry (Figure 5F). Briefly, TTC1-GFP HEK239T cells (as well as HEK293T cells exogenously expressing GFP alone as a control) were trypsinized, pelleted, washed with ice-cold 1x PBS and lysed in buffer containing 50mM HEPES pH 7.5, 200mM NaCl_2_, 2mM MgAc_2_, 1 mM DTT, 1% triton x-100 and 1x Pierce complete protease and phosphatase inhibitor cocktail (Thermo Fisher, A32959). Lysates were then clarified to remove debris by rotating end-over-end for 30 minutes at 4°C and centrifuging at 15,000xg for 10 minutes at 4°C. 50 ul of Pierce Streptavidin Magnetic Beads (Thermo Fisher, 88816) were first equilibrated with wash buffer (same composition as the lysis buffer except for containing 0.015% triton x-100).The beads were then immobilized with 16.67 ug biotinylated His14-Avi-SUMOEu1-tagged anti-GFP nanobody expressed and purified as previously described (Pleiner et. al, 2020) for 20 minutes on ice with occasional mixing. Remaining biotin binding sites were blocked by incubation with 100 uM biotin in 50mM HEPES pH 7.5 for 5 minutes on ice with occasional mixing. Beads were then washed and incubated with cleared cell lysate head-over-tail at 4°C for two hours (binding step). Post binding, beads were separated with a magnet, and 50 ul of each sample was saved as a flow-through fraction. The beads were washed 4 times with 1 ml wash buffer and bound material was eluted by cleavage with 250 nM SENPuB protease expressed and purified as previously described (Pleiner et. al, 2020) in minimal volume (20 ul) wash buffer for 30 minutes at 4°C with occasional mixing. Elutions were collected by separating the beads with a magnet and collecting the samples. Samples were flash frozen in liquid nitrogen and stored long term at -80°C. To visualize proteins, samples were first boiled with 6x Laemmli SDS buffer ar 95°C for 5 minutes and proteins were separated on Bolt 4-12% Bis-Tris gels (Thermo Fisher, NWO4122BOX and NWO4127BOX). Gels were washed in water, incubated with ReadyBlue Protein Gel Stain (Sigma RSB-1L) overnight for Coomassie staining, and destained in water. The gels were imaged on a Licor Odyssey CLx.

To determine relative HSP90 binding levels of wild-type and mutant exogenous GFP-tagged TTC1 expressed in HEK293T cells (Figure S9C), lysis was performed without detergent. Cells were first transduced with lentivirus carrying GFP-tagged wild-type and mutant exogenous TTC1 constructs (as well as a GFP-only control construct), sorted on the basis of GFP fluorescence and plated. After a day, cells were washed with ice-cold 1x PBS and then incubated with isolation buffer containing 10 mM HEPES pH 7.5, 225 mM sucrose, 75 mM mannitol, 0.1% (w/v) BSA and 1x Pierce complete protease and phosphatase inhibitor cocktail (Thermo Fisher, A32959). Cells were then scraped off the plate in isolation buffer, pelleted, and re-suspended in ice-cold hypotonic buffer containing 120 mOsm sucrose, 0.1% BSA, 10 mM HEPES ph 7.5 and 1x Pierce complete protease and phosphatase inhibitor cocktail for 5 mins. Cells were checked for swelling under a microscope, and then lysed with 20 strokes using a 26-gauge needle (BD PrecisionGlide, 305111) and a 1 ml syringe at 4°C. Hypertonic solution (1.25M sucrose in 10mM HEPES pH 7.5 buffer) was added to bring tonicity back up to 320 mOsm. Lysate was clarified to remove cell debris by centrifuging for 5 minutes at 930xg at 4°C. Streptavidin bead preparation, binding and elution was performed as previously described, with the wash buffer in this case being the isolation buffer. To detect changes in binding partners, samples were boiled in 6x Laemmli SDS buffer ar 95°C for 5 minutes, separated on Bolt 4-12% Bis-Tris gels and visualized by immunoblotting as described above.

### Protein expression and purification

The BL21(DE3) expression strain was used to express WT and mutant TTC1 in Luria-Bertani broth. Cultures were grown to an optical density of 1.0, induced with 1 mM IPTG, and then allowed to express at 18°C overnight. Cells were pelleted by centrifugation and the resulting pellets were resuspended in a buffer containing 500 mM NaCl, 50 mM Tris pH 7.5, 10 mM imidazole, 5 mM β-mercaptoethanol.

Prior to purification, reuspsnesions were supplemented with EDTA-free protease inhibitor tablets (Roche) and lysozyme, and then subsequently lysed by sonication. Lysate was then centrifuged at 18,000 rpm for 30 minutes in an SS-34 rotor. The supernatant was then incubated with NiNTA (Qiagen, Valencia, CA) resin for 30 minutes while rolling at 4°C. NiNTA resin was then washed with 20x column volumes of resuspension buffer, and then with 5x column volumes of protease elution buffer containing 150 mM NaCl, 50 mM Tris pH 7.5, 10 mM imidazole, 5 mM β-mercaptoethanol, and 10% glycerol. NiNTA resin was then incubated with bdSENP1 for 2 hours at 4°C to release SUMO-cleaved protein from the resin. Next, cleaved protein was concentrated and injected onto a Superdex 200 increase 10/300 GL size exclusion column equilibrated with buffer containing 150 mM KAcetate, 50 mM HEPES pH 7.4, 2 mM MgAcetate, and 1 mM DTT. Peak fractions were analyzed by SDS-PAGE, and fractions containing full length protein were pooled, concentrated, aliquoted, and flash frozen.

### PURE system solubilization assay

Purified CaM or GFP-TTC1 (WT or mutant) were added to PURE translations reactions of His-CISD1-Flag at 12 uM. 13.5 ul reactions, incubated at 32°C for 2 hours, were size fractionated on a 200 ul 5-25% sucrose gradient in PSB (supplemented with 100 nM CaCl2 in the case of CaM complexes). To separate fractions, samples were allowed to sit on ice for 1 hour, and then spun in a TLS-55 rotor at 4°C for 120 min with the slowest acceleration and deceleration settings. Eleven 20 ul fractions were collected from the top, and protein sample buffer was added directly for SDS-Page analysis.

### In vitro translation and mitochondrial insertion

Mitochondria for insertion experiments was isolated as previously described (Guna et. al, 2022). In vitro translations for insertion reactions were done using rabbit reticulocyte lysate (Guna et. al, 2018). Templates for PCR were based on the SP64 vector (Promega, USA), and amplified by primers binding upstream of the SP64 promoter and roughly 200 bp downstream of the stop codon (Sharma et. al, 2010) for use in transcription reactions. Following transcription at 37°C for 1.5 hours, reactions were directly used to translate substrate in RRL for 15-30 min at 32°C in the presence of radioactive 35S-Methionine. For assessing engagement by TTC1 in RRL (Figure 6A), substrates were released from the ribosome by incubation with 1mM puromycin at 32°C for 10 minutes followed by immunoprecipitation with packed FLAG beads (90 minutes at 4°C) and separation of bound proteins by SDS-PAGE as previously described. For assessing the ability of TTC1 to promote insertion into isolated mitochondria (Figure 6E), purified wild type or mutant TTC1 was added at increasing concentrations (0.5, 1, 2.5, or 5 uM). Insertion reactions were performed by adding 4 ul of translation reaction into 50 ul of mitochondrial import buffer (250 mM sucrose, 5 mM Mg(Ac)2, 80 mM KAc, 20 mM HEPEs pH 7.4, 2.5 mM succinate) with 15 ug of purified mitochondria and further incubated at 32°C for 30 min.

Protease digestion to assess substrate insertion was identical to previous studies (Guna et. al, 2022). Briefly, proteinase K was added at 0.25 mg/ml, and reactions were incubated on ice for 1 hour. Reactions were then quenched by 5 mM PMSF in DMSO and transferred to boiling 1% SDS in 0.1M Tris/HCl pH 8.0. Immunoprecitation for His-tagged elements was done by incubation of PK treated reactions with NiNTA resin in IP buffer (50 mM HEPES pH 7.5, 500 mM NaCl, 10 mM imidazole, 1% Triton). PK treated reactions were diluted to 1 ml in buffer and mixed with 20 uL resin, with end-to-end mixing for 1.5 hours at 4°C. Following three washes with 1 x 1 ml IP buffer, products were eluted in sample buffer containing 50 mM EDTA at pH 8.0.

### Quantification and statistical analysis

### Clustering analysis and minimal distortion embedding

All clustermaps were generated using the Python seaborn library clustermap function with the average linkage UPGMA (Unweighted Pair Group Method with Arithmetic Mean) method computing Euclidean distances. To generate the clustermap in Figure S3, hierarchical clustering was performed on all raw fold changes in GFP:RFP ratios for the respective perturbations and reporters. For the factor-factor and reporter-reporter clustermaps in Figure. 2, correlation matrices were generated using the pandas library before hierarchical clustering in the same manner as described above. Minimal distortion embedding to visualize the correlation of TTC1 with other genes in genome-scale perturb-seq data was performed as previously described (Replogle and Saunders et. al, 2022).

### Imaging colocalization analyses

Quantification of 3D-reconstructed z-stacks of images taken on the Zeiss LSM980 microscope for endogenous substrate localization analysis in Figures 4 and S6 was performed using Oxford’s Imaris analysis software. For all conditions, 15-20 images were taken at 1.5x magnification, each containing ∼5-8 individual cells in a single field of view, with a total of ∼100 cells analyzed across each condition. First, surfaces were assigned to each channel (DAPI, ER, MitoTracker and Substrate) using the Imaris Surface algorithm. Images were batch processed across all conditions (Control, NACA and TOMM70 knockdown; Figures 4 and S6) to consistently construct surfaces for all channels using the Imaris Batch package. Percent co-localization was determined by selecting ‘shortest distance to surface’ under the Imaris ‘filter’ option, setting the shortest distance to 0 to select overlapping signal from channels (ex. from substrate to ER, substrate to mitochondria and ER to mitochondria). To determine percent co-localization of a substrate to ‘ER/mitochondria overlap’ vs ‘ER alone’ surfaces as shown in Figures 4 and S6, the analysis was performed in two steps: first, the ER/mitochondria overlapping surface was constructed using the surface algorithm and ‘shortest distance to surface’ filter (from ER to mitochondria) as described above. Then, the surface for ‘percent substrate co-localized to ER total’ (calculated as described above) was co-localized to the new ‘ER/mitochondria’ surface, using the same algorithms and selection filters as previously described. Finally, the percent co-localized to ‘ER alone’ was calculated by subtracting the percent co-localized to ‘ER/mitochondria’ from the percent co-localized to ‘ER total’. For all pairwise statistical comparisons, means with the standard deviation are plotted. P-values are calculated using unpaired t tests with Welch’s correction. For plots with multiple comparisons, p-values were adjusted using the Holm-Sidak method with Prism Graphpad. Significance values were all set to 0.05.

### Imaging correlation analyses

For the images shown in Figures 4 and S6, correlation analyses were performed using the ImageJ JaCoP colocalization module as previously described (Bolte and Coredelières, 2006). All correlation values plotted are Pearson correlation coefficients. For all pairwise statistical comparisons, means with the standard deviation are plotted. P-values are calculated using unpaired t-tests with Welch’s correction. For plots with multiple comparisons, p-values were adjusted using the Holm-Sidak method with Prism Graphpad. Significance values (alpha) were all set to 0.05.

### Imaging fragmentation analyses

Quantification of 3D-reconstructed z-stacks of images taken on the Zeiss LSM980 microscope for mitochondrial and ER network analysis in Figures 5, S8 and S9 was performed using Oxford’s Imaris analysis software. For all conditions, 10 images were taken at 1.5x magnification, each containing ∼5 individual cells in a single field of view, with a total of ∼50 cells analyzed across each condition. In each case, the mitochondrial and ER surfaces were assigned used the Imaris surface algorithm and images were subsequently batch-processed. To calculate the extent of network fragmentation, the ‘count/area’ metric was calculated – total count (number of individual ‘pieces’ assigned to the organelle surface, rather than a continuous network) and total area were measured using the surface algorithm again. A higher count/total area indicates a greater extent of fragmentation. For mitochondrial network analysis, given the significant disparity in absolute values between control and TTC1 kd cells, log10 (extent of fragmentation) values are plotted and the same was done for ER network analysis for consistency. P-values are calculated using unpaired t tests with Welch’s correction. For datasets comparing three or more sets of unpaired measurements (Figures S8 and S10), the Welch and Brown-Forsythe ANOVA test was used in Prism Graphpad (assumed to be sampled from a Gaussian distribution without equal variances). Significance values (alpha) were all set to 0.05.

### Statistics

All data are shown with means and SD plotted. For Seahorse experiment analysis, the Welch and Brown-Forsythe ANOVA test was used in Prism Graphpad (assumed to be sampled from a Gaussian distribution but without equal variances). Significance values were all set to 0.05. Experiment specific details can be found in respective figure legends.

## SUPPLEMENTARY MATERIALS

Figures S1-S10

Tables S1-S10

### SUPPLEMENTARY FIGURE LEGENDS

**Figure S1. Characterization of α-helical reporter integration in the context of CRISPRi screens (related to Figure 1)**

(A) The localization of reporters used for genome-wide CRISPRi screens (Figure 1A) was assessed using microscopy. Indicated GFP11-fused α-helical proteins were expressed in RPE1 ZIM3 CRISPRi cells constitutively expressing IMS GFP1-10 and stained with MitoTracker. Merge colors: Green: GFP, Magenta: MitoTracker. (B) CISD1, FUNDC1 and TSPO were conjugated to GFP11 at either the N- and C-termini, expressed in K562 IMS GFP1-10 cells, and analyzed by flow cytometry with the GFP:RFP ratio plotted. All three reporters show insertion in the correct, predicted topology into the mitochondrial outer membrane. (C) Monoclonal K562 CRISPRi cell lines were constructed for screening with constitutive expression of IMS-targeted GFP1-10 and tet-inducible expression of the indicated GFP11-fused RFP-P2A reporter (Figure 1A). Cells were transduced with a genome-scale CRISPRi sgRNA library, subjected to puromycin selection, and then induced with doxycycline 36 hours prior to sorting. Cells were then sorted based on the ratiometric change of GFP relative to RFP, indicative of stable reporter integration in the outer membrane. sgRNAs expressed in the sorted cells were isolated and identified by deep sequencing.

**Figure S2. Comparison of relevant hits across genome-wide CRISPRi screens (related to Figure 1)**

(A) Plots comparing the average phenotype scores (based on GFP:RFP ratios) of genes of interest across CRISPRi genome-wide screens with the same color scheme as in Fig 1B. For SA protein CISD1 which is expected to regulate iron sulfur cluster biogenesis, partner factors such as NARFL, FAM96A, NFS1 and ABCB7 are highlighted in dark gray, which all have enzymatic roles in iron sulfur cluster biogenesis. Additionally, TOMM20 and TOMM22 are highlighted in orange as factors that show consistent destabilizing effects across all reporter screening conditions. (B) Using our ratiometric fluorescent reporter system, we demonstrate that CRISPRi knockdown of TOMM20 or (C) TOMM22 with two independent guides leads to a loss of complementation of GFP11 fused to either the MICU1 or LACTB targeting signal relative to the RFP expression control in K562 IMS GFP1-10 cells. Similarly, we also observed a pronounced decrease in complementation of GFP11-fused TIMM9A, a canonical IMS substrate. The extent of this decrease in complementation is similar to that seen for our screen reporters (FUNDC1 and OMP25 are shown here). Together, these data suggest that TOMM20/TOMM22 knockdown screen phenotypes are a result of a more general effect on GFP1-10 levels in the IMS.

**Figure S3. Comprehensive arrayed analysis of the dependencies of α-helical mitochondrial outer membrane proteins on putative biogenesis factors (related to Figure 2)**

(A) Schematic of the arrayed screen used to systematically test the dependencies of a diverse panel of α-helical outer membrane reporters on putative biogenesis and quality control factors. K562 ZIM3 CRISPRi cells constitutively expressing IMS GFP1-10 were first transduced with guides targeting each factor of interest or a non-targeting control sgRNA. Respective reporters were expressed and analyzed by flow cytometry and a GFP:RFP ratio was calculated. Two independent experiments were performed. The resulting heat map represents 9 factors of interest (and one non-targeting control guide) and 21 reporters and is colored by the fold change in the GFP:RFP ratio for factor knockdown normalized to the control, non-targeting condition. Hierarchical clustering on the raw fold changes initially organizes factors and reporters into putative biogenesis pathways, with reporters largely organized by topology. (B) Integration of indicated GFP11-fused reporters with TOMM70A and MARCHF5 depleted relative to the non-targeting control in K562 IMS GFP1-10 cells. The unique dependencies of TSPO on both factors explains its separation from two other polytopic proteins FUNDC1 and MTCH2 in the clustering. (C) Knockdown of each indicated factor across both replicates was measured using qRT-PCR normalized to the housekeeping gene GAPDH (see methods for details).

**Figure S4. Patterns of correlation and hierarchical clustering are strongly correlated across replicates (related to Figure 2)**

(A) Correlation values for each factor-factor pair are plotted across arrayed screen replicates. Factor pairs of interest are highlighted in blue. (B) Similarly, correlation values for each reporter-reporter pair are plotted across replicates. Reporter pairs of interest are highlighted based on their topologies.

**Figure S5. Proteomic and transcriptional changes under combined knock-down of putative biogenesis factors (related to Figure 3)**

(A) (Left) Immunoblotting for TTC1, MTCH2 and a loading control (vinculin) for K562 ZIM3 CRISPRi IMS GFP1-10 cells expressing sgRNAs targeting either i) TTC1 and MTCH2 (using a dual guide), ii) TTC1, iii) MTCH2, iv) non-targeting control sgRNA. (Right) qRT-PCR was used to assess expression of the indicated genes normalized to housekeeping gene GAPDH for the same conditions. (B) Immunoblotting for TOMM70, MTCH2 and loading control (tubulin) for cells expressing sgRNAs targeting the indicated genes (i) TOMM70A and MTCH2 with a dual guide, ii) TOMM70A, iii) MTCH2, iv) non-targeting control). (C) TTC1, MTCH2 and TOMM70 levels across the various conditions indicated in Figures 3E and 3G are assessed using immunoblotting with vinculin as the loading control.

**Figure S6. NACα depletion phenocopies the effect of TOMM70 depletion on endogenous TSPO localization (related to Figure 4)**

(A) The integration of GFP11-fused SA, TA, polytopic and IMS control reporters was assessed in K562 KOX1 and ZIM3 CRISPRi cells constitutively expressing IMS GFP1-10 and either a sgRNA targeting BTF3 (NACβ) or a non-targeting control sgRNA, as in Fig 4A. (B) To assess the effects of NACα depletion on mitochondrial signal-sequences, a MICU1(signal sequence)-GFP construct was expressed in RPE1 ZIM3 CRISPRi cells constitutively expressing either a sgRNA targeting NACα or a non-targeting control sgRNA and tracked using live-cell super-resolution confocal microscopy. Merge colors: Green: GFP, Magenta: MitoTracker. (C) Magnification of endogenous TSPO signal under NACα and TOMM70A depletion relative to the non-targeting control from Figure 4B shows similar mis-localization and punctate appearance. (D) Quantification of endogenous TSPO and MAVS localized to ER/MitoTracker overlapping and solely ER surfaces for the images in 4B. This allows us to specifically track the extent of mis-localization of substrates to the ER outside of the mitochondria under NACα depletion. (E) Comparison of percent co-localization of endogenous TSPO to the mitochondria and ER between the NACα and TOMM70A depleted cells from 4B. (F) In an orthogonal method of analysis, correlation between endogenous TSPO and the mitochondria, endogenous TSPO and the ER, and the ER and mitochondria for the images in 4B was assessed using the ImageJ JACoP plugin. Pearson correlation values are reported. For (D)-(F), error bars show mean ± SD of 100 cells. Statistical significance was evaluated by multiple unpaired t-tests with the Holm-Sidak multiple test correction. *, p < 0.05. **, p < 0.01. ****, p < 0.0001. ns (non-significant), p > 0.05.

**Figure S7. Point mutations in the ribosome-binding domains of NACα and NACβ affect integration of outer membrane α-helical reporters (related to Figure 4)**

(A) Flow data from the heatmap in Figure 4D for two polytopic reporters FUNDC1 and MTCH2 normalized based on the GFP:RFP ratios and plotted as cdf plots. (B) NACα and NACβ levels across the various conditions in Figure 4D were assessed using immunoblotting along with a loading control (vinculin). (C) qRT-PCR used to assess gene expression of NACα and NACβ for KOX1 and ZIM3 CRISPRi relative to the housekeeping gene GAPDH.

**Figure S8. TTC1 clusters with genes known to impact mitochondrial biogenesis and specifically affects the mitochondrial outer membrane (related to Figure 5)**

(A) Minimum distortion embedding for all essential genes in K562 cells, where each dot represents a genetic perturbation from genome-scale perturb-seq data (Replogle and Saunders et. al, 2022) that places genes with correlated expression profiles nearby (see Methods). A cluster with TTC1 is expanded to show neighbors involved in mitochondrial biogenesis and function. (B) Integrated stress response (ISR) and Unfolded protein response (UPR) scores for perturbation of all essential genes in K562 cells are measured as previously described (Replogle and Saunders et. al, 2022) using the sum of z-normalized expression of ISR or UPR marker genes and plotted against each other (higher score indicates stronger activation of the pathway). TTC1 is highlighted in gold, ISR activators and mitochondrial biogenesis genes in purple, and UPR activators in turquoise. (C) Integration of inner membrane (Cox4) and matrix (HSP60) GFP11-fused reporters in K562 IMS and Matrix GFP1-10 cell lines respectively under TTC1 depletion. (D) Similarly, integration of ER signal-sequence (CalR), tail-anchored (SQS, VAMP2, Sec61β, RHOT1) and polytopic (ACER1) GFP11-fused reporters in K562 ER GFP1-10 cells expressing sgRNAs targeting i) TTC1, ii) EMC2 (ER biogenesis factor), iii) ATP13A1 (ER quality control factor) was evaluated relative to cells expressing non-targeting sgRNAs (control). Data is represented as a heatmap colored by the fold change in reporter integration (GFP:RFP ratio) for each condition relative to the control. Flow data for the SQS-GFP11 reporter under all indicated conditions is normalized based on the GFP:RFP ratio and plotted as histograms. (E) Knockdown of the genes in (D) is assessed using qRT-PCR relative to the housekeeping gene GAPDH. (F) Super-resolution confocal microscopy showing mitochondrial fragmentation for RPE1 ZIM3 CRISPRi cells expressing a sgRNA against TTC1 and either i) exogenous TTC1 or ii) exogenous MTCH2. (G) Quantification of the morphology shown in (F) compared to control and TTC1 knockdown mitochondrial morphology from Figure 5A. Error bars show mean ± SD of 50 cells. Statistical significance was evaluated using the Welch and Brown-Forsythe ANOVA test. *, p < 0.05. **, p < 0.01. (H) Relative mRNA levels of outer membrane proteins depleted in TTC1 knockdown cells (Figures 5D and 5E) assessed using qRT-PCR (relative to the housekeeping gene GAPDH) show that the substrate biogenesis defects are largely post-transcriptional.

**Figure S9. TTC1 association with HSP90 promotes efficient SA protein biogenesis in vivo (related to Figure 6)**

(A) AlphaFold2 complete model of TTC1 colored by evolutionary conservation as indicated in Figure 6B, with the N-terminal loop pLDDT score noted. (B) Expression of exogenous WT and mutant TTC1 rescue constructs used in Figure 6C was assessed by immunoblotting with vinculin as the loading control. (C) (Left) Exogenous GFP-tagged WT and TPR mutant (R190A + R194A) constructs were expressed in HEK293T cells, purified using an anti-GFP nanobody under native conditions, and HSP90 binding was assessed by immunoblotting. (Right) HSP90 binding of GFP-tagged C-terminal I271D mutant TTC1 was assessed in the same manner with GFP transduction and immunoprecipitation used a control for non-specific binding in both cases.

**Figure S10. The TTC1 C-terminal I271D mutant can partially rescue mitochondrial phenotypes (related to Figure 6)**

A) (Top) Super-resolution confocal microscopy assessing mitochondrial morphology for RPE1 ZIM3 cells constitutively expressing a sgRNA targeting TTC1 and either i) no exogenous TTC1, ii) exogenous WT TTC1 or iii) exogenous I271D-TTC1 compared to cells expressing a non-targeting sgRNA (control). White arrows indicate mitochondrial fragmentation. (Bottom) Quantification of mitochondrial morphology as described in Figures 5A and S8H. Error bars show mean ± SD of 50 cells. Statistical significance was evaluated using the Welch and Brown-Forsythe ANOVA test. **, p < 0.01, ***, p < 0.001. (B) Extent of rescue of WT and C-terminal I271D mutant TTC1 constructs on mitochondrial respiration using a Seahorse assay in K562 cells as described in Figure 5C. Data are presented as average ± SD, n = 10. (C) Individual parameters from (B) such as basal respiration, spare respiratory capacity, maximal respiration, coupling efficiency and ATP production are separately plotted across all conditions. Error bars show mean ± SD of 10 wells with 4x10^4^ cells each. Statistical significance was evaluated using the Welch and Brown-Forsythe ANOVA test. *, p < 0.05. **, p < 0.01, ****, p < 0.0001. ns (non-significant), p > 0.05. (D) Levels of TTC1 across the conditions in Figures 5C and S9C were assessed by immunoblotting with vinculin as the loading control. (E) Purifications of TTC1 constructs from *E.Coli* cells used for in vitro experiments in Figures 6D and 6E.

### SUPPLEMENTARY DATA TABLES

**Table S1 (related to Figure 1): CISD1 Screen.** Genome-wide CRISPRi screen data table for all genes perturbed (Figure 1B)

**Table S2 (related to Figure 1)**: **FUNDC1 Screen**. Genome-wide CRISPRi screen data table for all genes perturbed (Figure 1B)

**Table S3 (related to Figure 1)**: **TSPO Screen**. Genome-wide CRISPRi screen data table for all genes perturbed (Figure 1B)

**Table S4 (related to Figure 1)**: **OMP25 Screen**. Genome-wide CRISPRi screen data table for all genes perturbed (Figure 1B)

**Table S5 (related to Figure 2): Arrayed CRISPRi Screen**. Data table with normalized GFP:RFP ratios (to control sgRNA) for all perturbations and all reporters in duplicate (Figures 2 and S3)

**Table S6 (related to Figure 5): TTC1 knock-down TMT mass spectrometry data**. Complete data table. Sheet 1: Raw quantification from whole-cell extracts. Sheet 2: Data normalized to the mitochondrial proteome as defined by MitoCarta 3.0 (Figure 5D)

**Table S7 (related to Figures 2, 3, 4, 5 and 6): Microscopy and qRT-PCR individual data values**. Sheet 1: Colocalization analyses using the Imaris software algorithms (see Methods for details). Each data point is from a single image, with ∼5-8 cells each for a total of 100 cells assayed across all conditions (Figures 4B and S6C-F). Sheet 2: Correlation analyses using ImageJ JaCoP. Each data point is from a single image as described for colocalization analyses (Figures 4 and S6). Sheet 3: Mitochondrial morphology analyses using Imaris software algorithms (see Methods for details). Each data point is from a single image, with ∼5-8 cells each for a total of 50 cells assayed across all conditions (Figures 5A, S8F-G, S10A). Sheet 4: ER morphology analyses using Imaris software algorithms (see Methods for details). Each data point is from a single image, with ∼5-8 cells each for a total of 50 cells assayed across all conditions (Figure 5B). Sheet 5: qRT-PCR data associated with Figures S3C, S5A, S7C, S8E and S8H.

**Table S8 (related to Methods): Plasmid spreadsheet**. All plasmids cloned and used in the study are detailed here.

**Table S9 (related to Methods)**: **Reagents spreadsheet**. Antibodies, chemical reagents and deposited data used in the study are detailed here.

**Table S10 (related to Methods)**: **Oligonucleotides spreadsheet.** All oligonucleotides used (qRT-PCR and sequencing primers) are detailed here.

