## Supplementary figures and images for "Triaging of ⍺-helical proteins to the mitochondrial outer membrane by distinct chaperone machinery based on substrate topology"

### Supplemental Figure 1

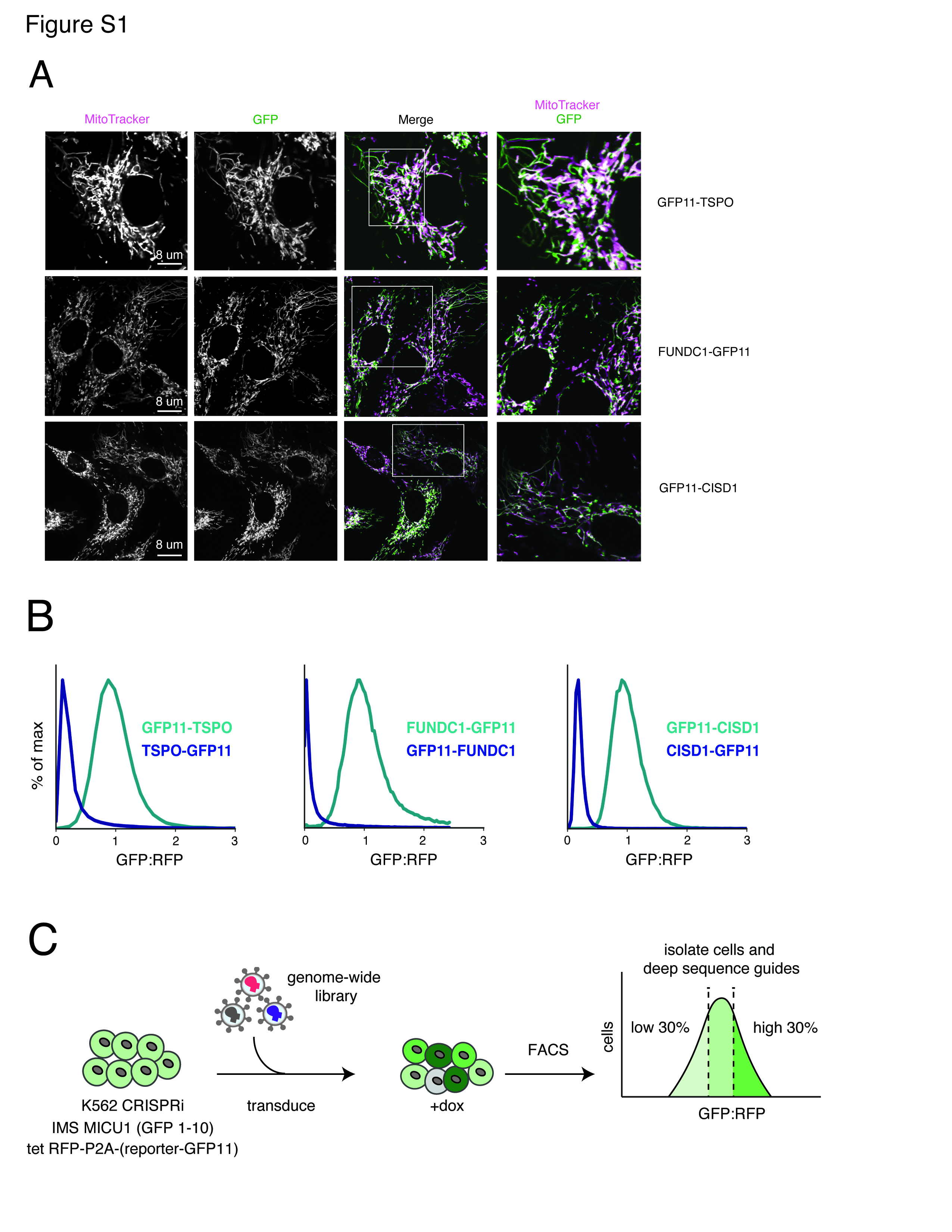

### Supplemental Figure 2

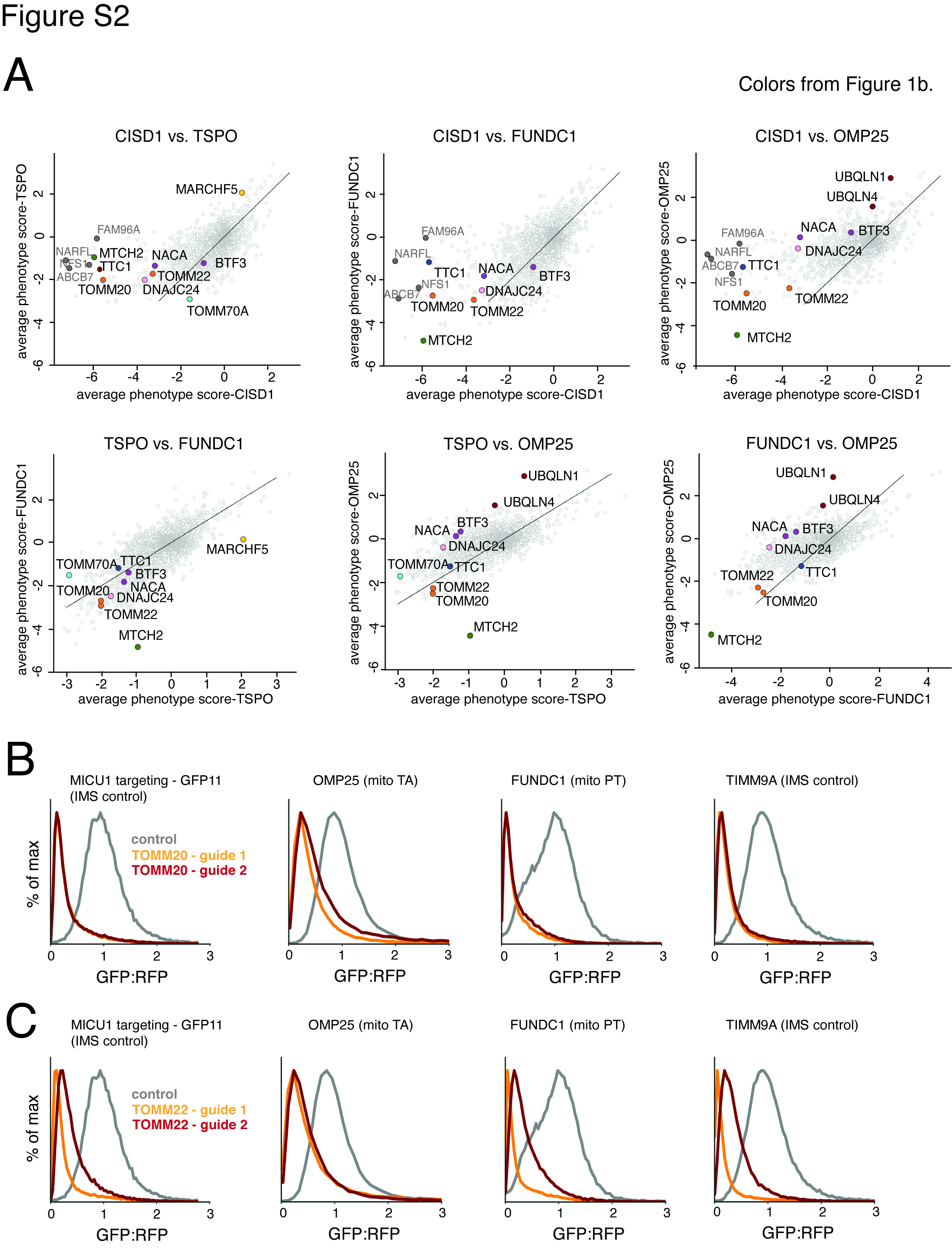

### Supplemental Figure 3

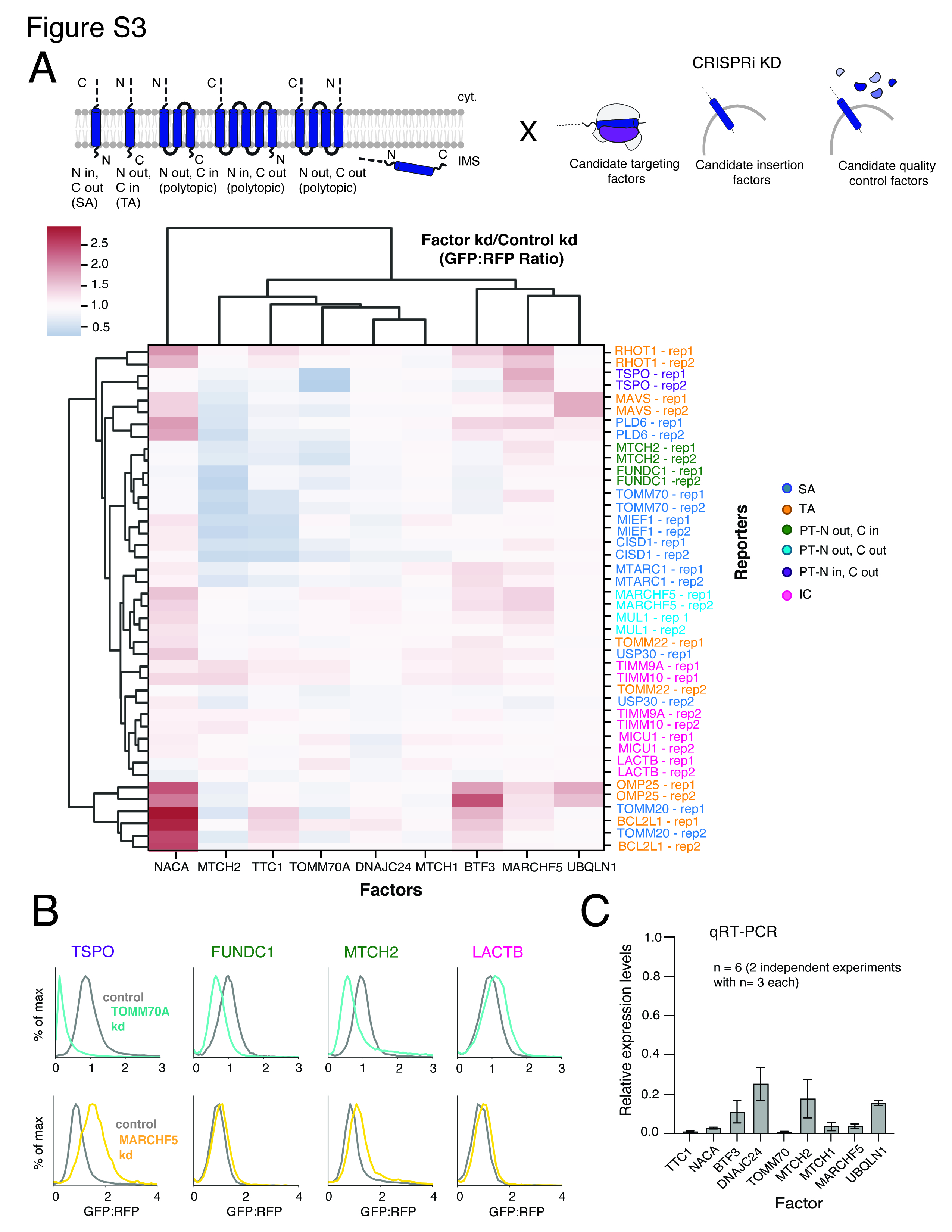

### Supplemental Figure 4

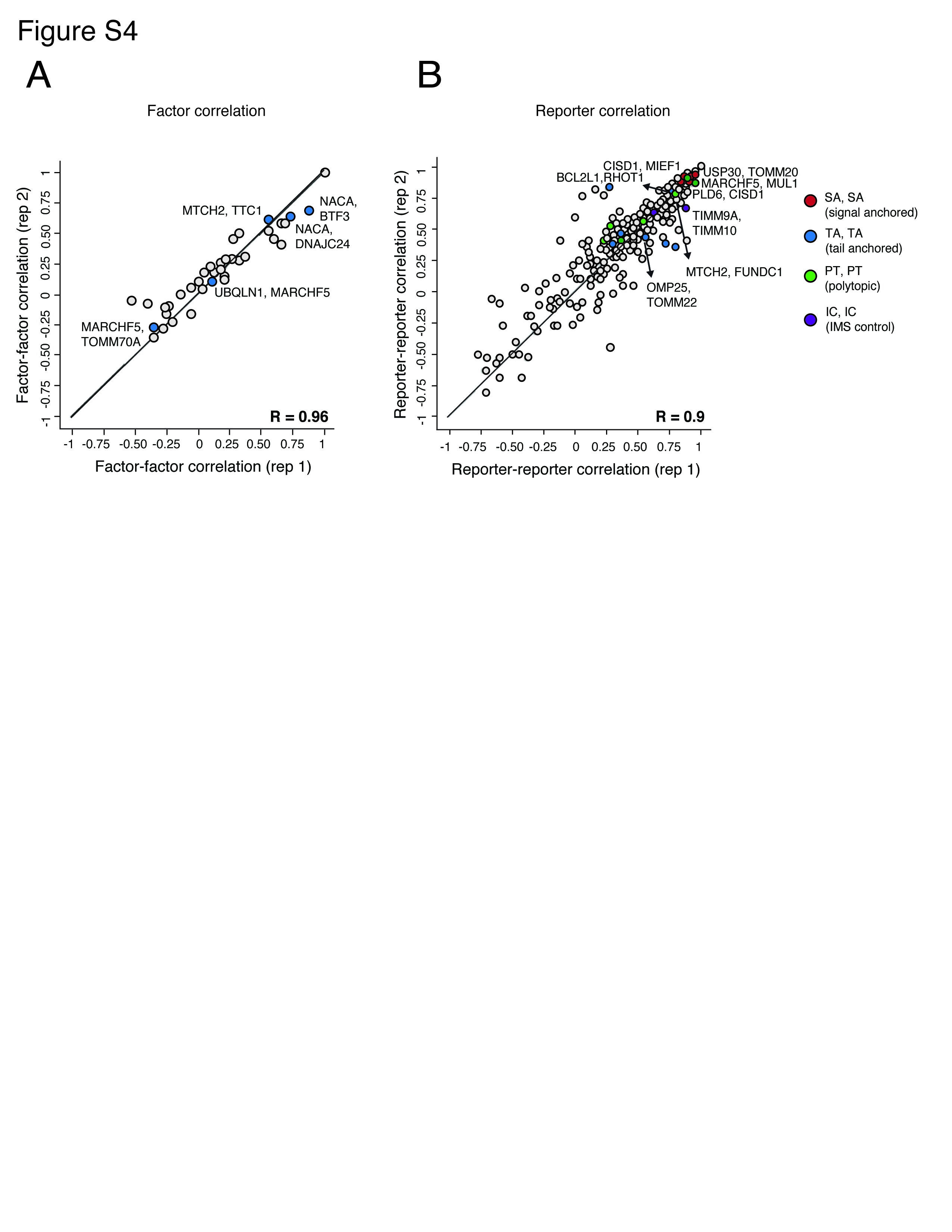

### Supplemental Figure 5

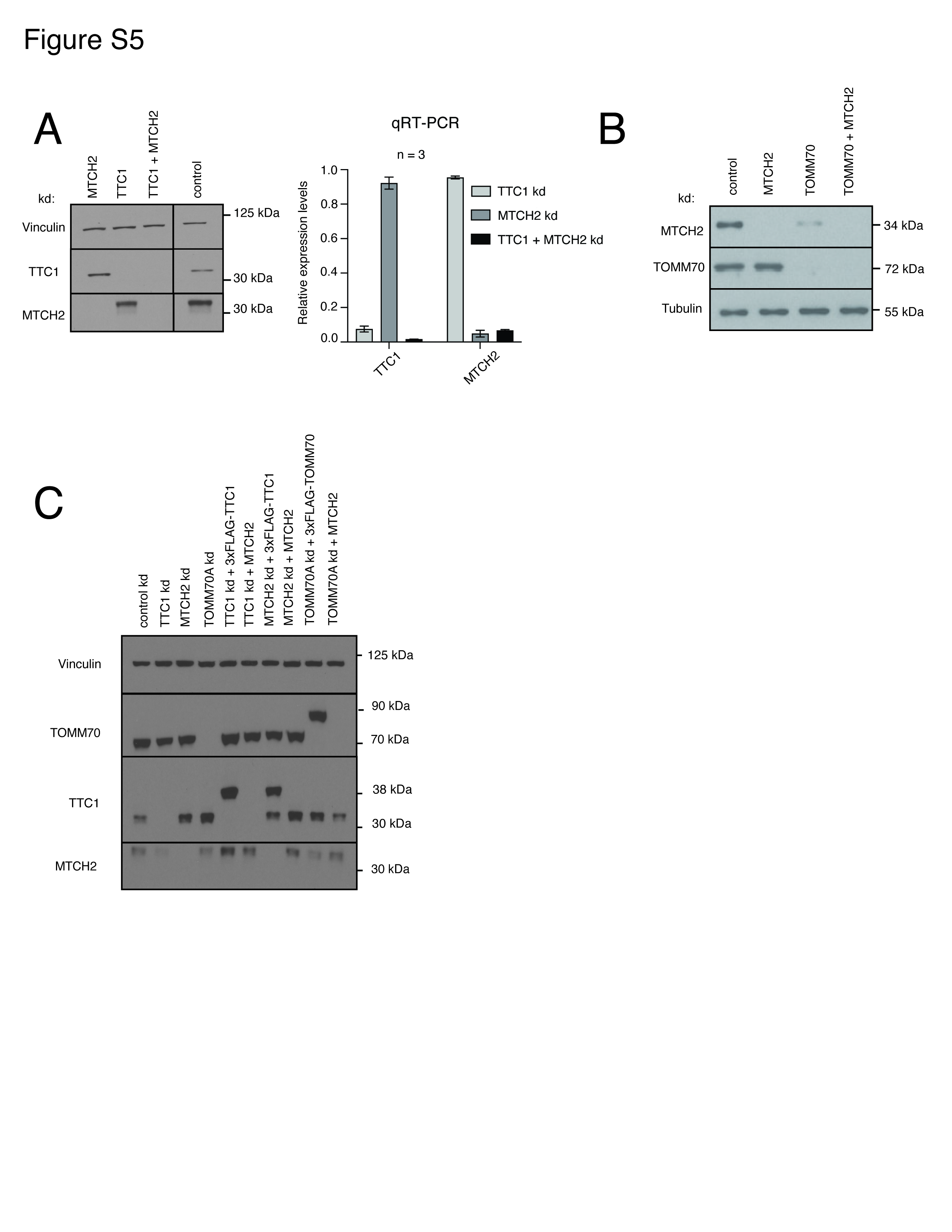

### Supplemental Figure 6

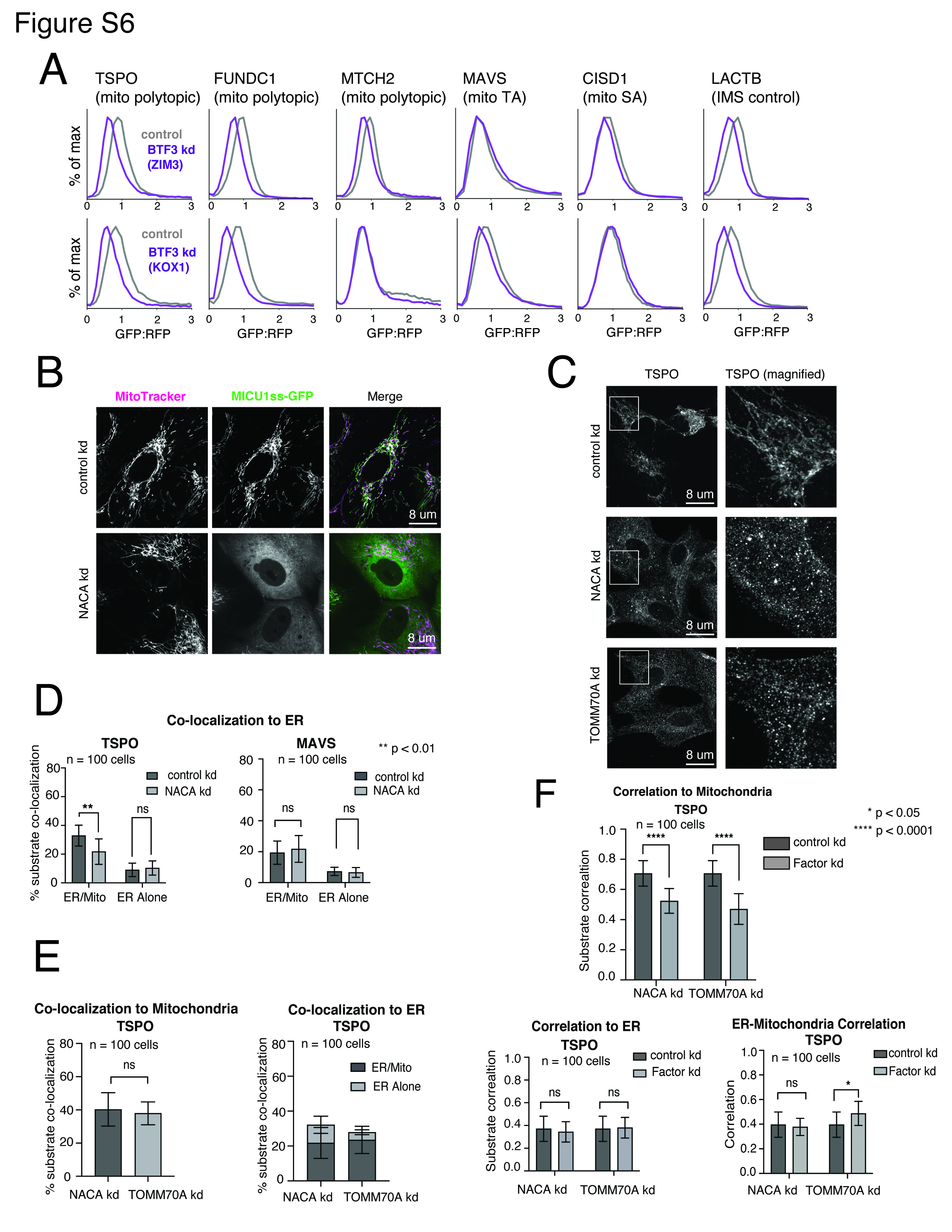

### Supplemental Figure 7

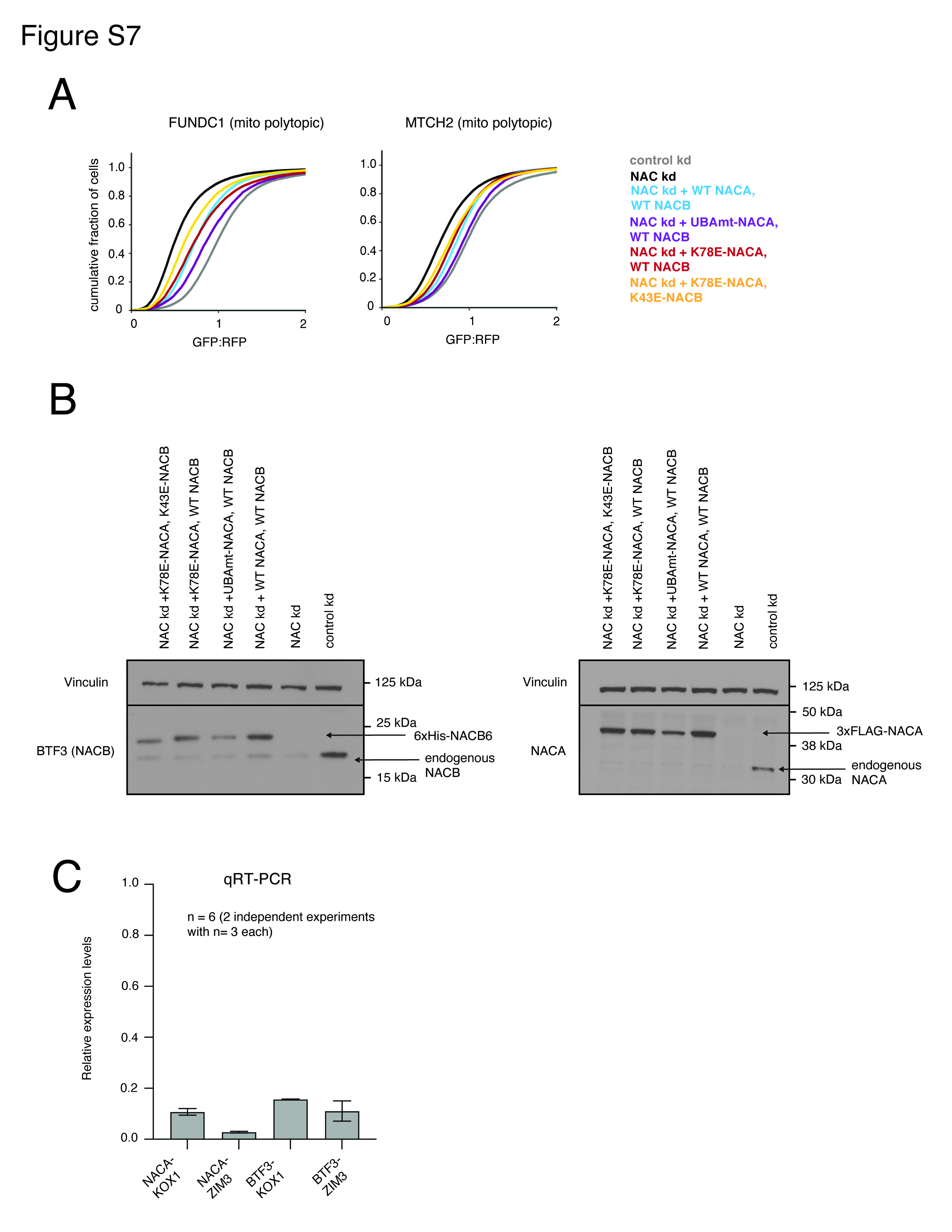

### Supplemental Figure 8

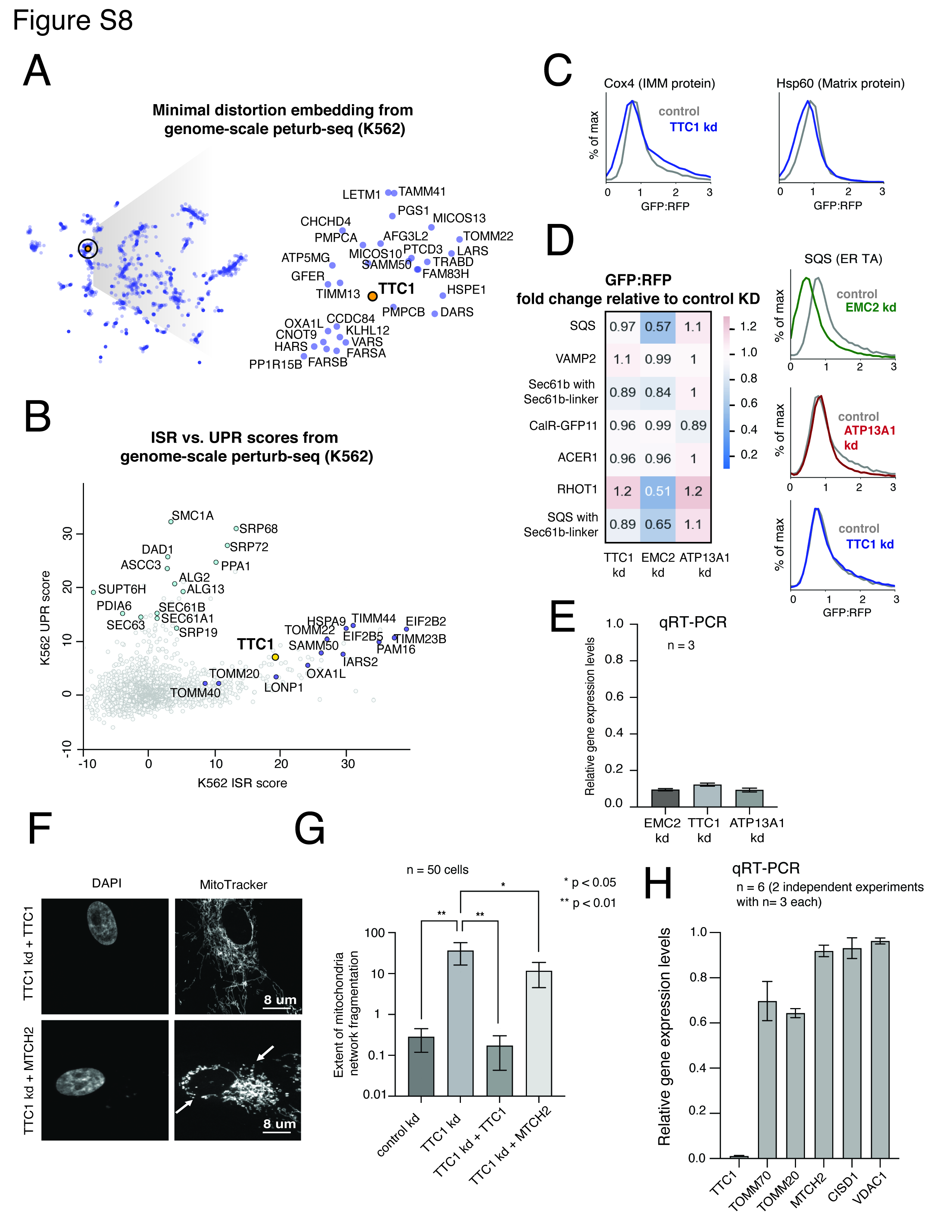

### Supplemental Figure 9

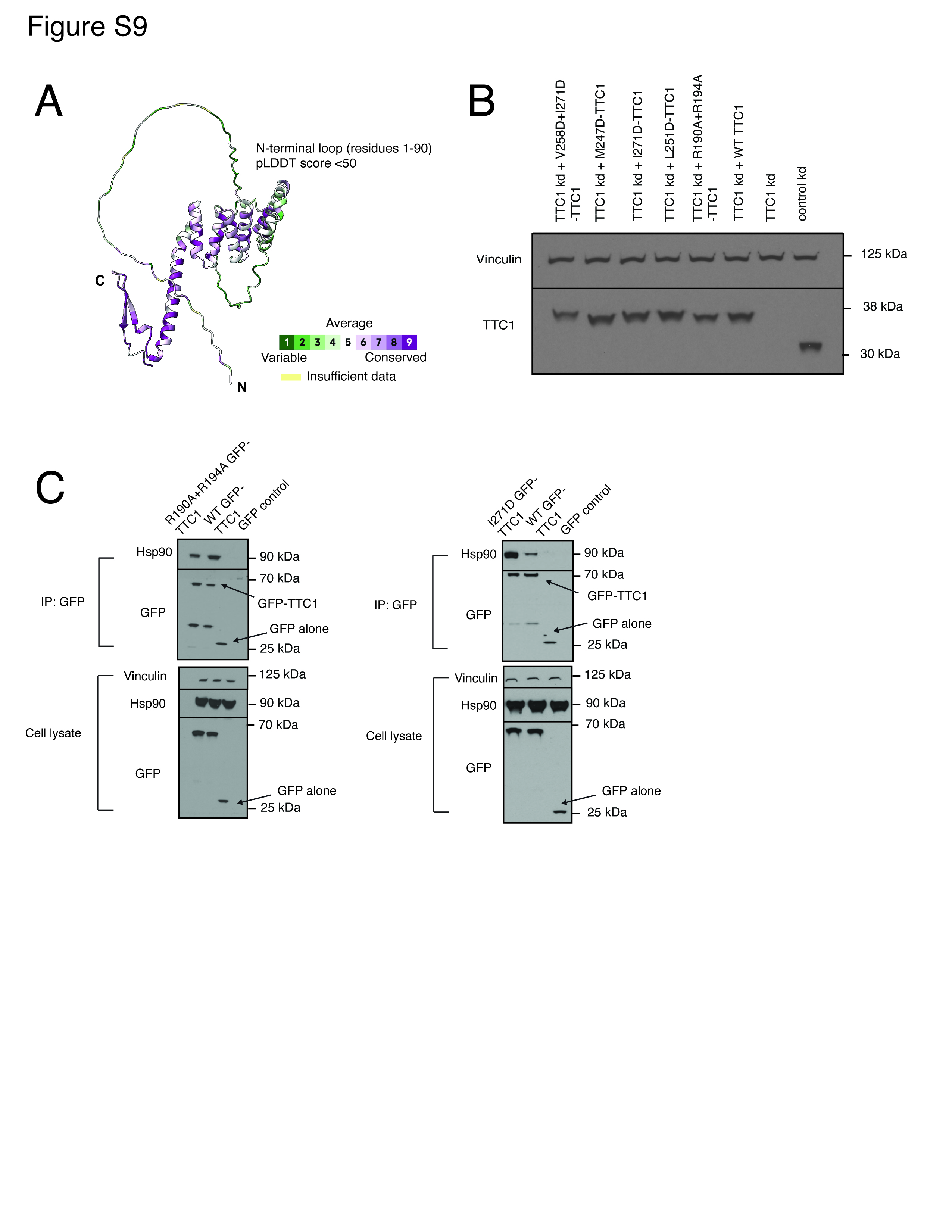

### Supplemental Figure 10

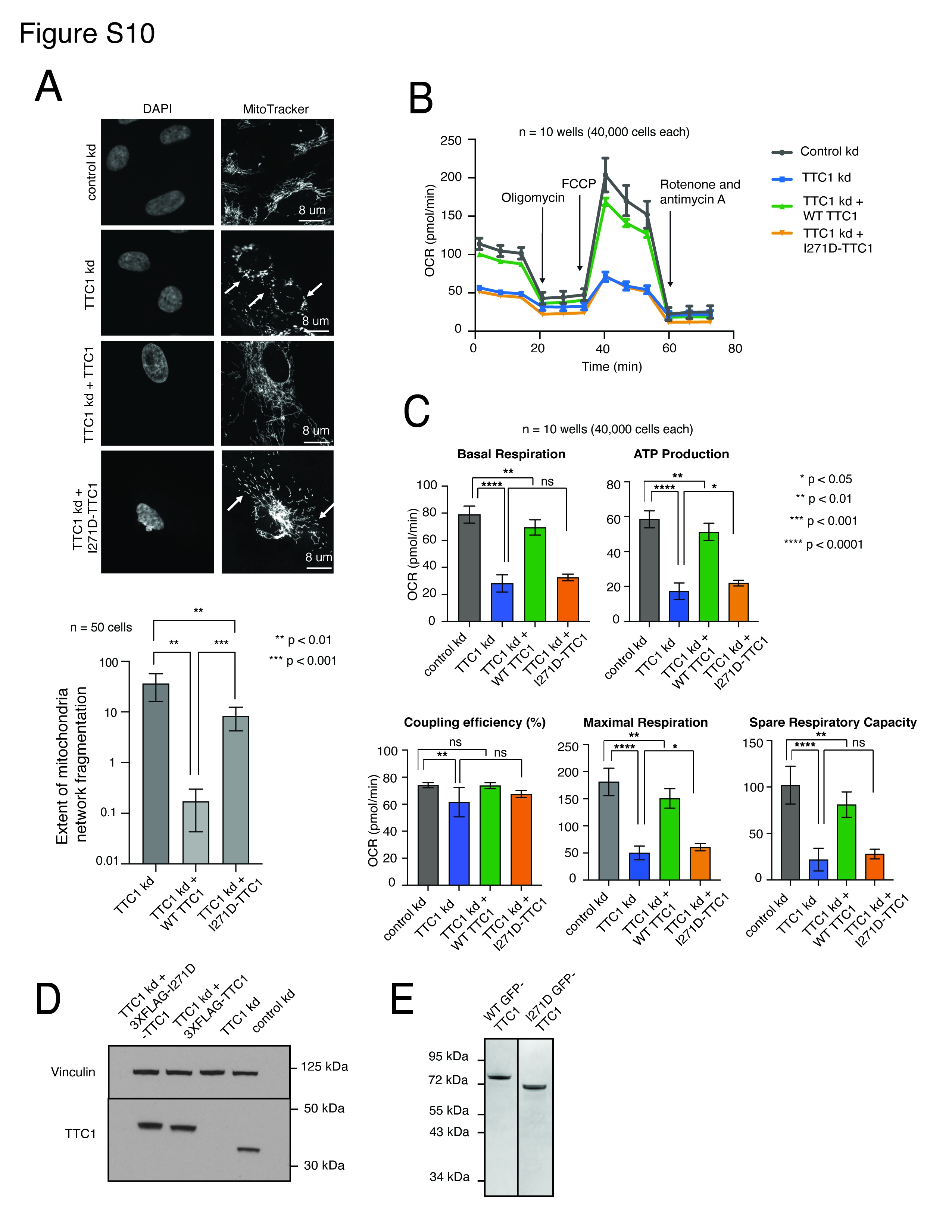
